# Targeting CISH enhances natural cytotoxicity receptor signaling and reduces NK cell exhaustion to improve solid tumor immunity

**DOI:** 10.1101/2021.03.16.435571

**Authors:** P-L. Bernard, R. B. Delconte, S. Pastor, V. Laletin, C. Costa Da Silva, A. Goubard, E. Josselin, R. Castellano, A. Krug, J. Vernerey, R. Devillier, D. Olive, E. Verhoeyen, E. Vivier, N. D. Huntington, J. A. Nunès, G. Guittard

## Abstract

Cytokine inducible SH2-containing protein (CISH) is a natural killer (NK) cell negative regulator of cytokine signaling pathway. To further understand CISH functions in NK cells, we developed a conditional *Cish*-deficient mouse model in NK cells (*Cish^fl/fl^Ncr1^Ki/+^*). We detected no developmental or homeostatic difference in NK cells.

However, global gene expression of *Cish^fl/fl^Ncr1^Ki/+^* NK cells compared to *Cish^+/+^Ncr1^Ki/+^* NK cells revealed upregulation of pathways and genes associated with NK cell cycling and activation. We show that CISH does not only regulate interleukin-15 (IL-15) signaling pathways but also natural cytotoxicity receptors (NCR) pathways. Indeed, CISH protein expression level increases upon NCR triggering. Primed *Cish^fl/fl^Ncr1^Ki/+^* NK cells display increased activation upon NCR stimulation. *Cish^fl/fl^Ncr1^Ki/+^* NK cells display lower activation thresholds and *Cish^fl/fl^Ncr1^Ki/+^* mice are more resistant to tumor metastasis. Remarkably, we found that *Cish^fl/fl^Ncr1^Ki/+^* mice were also more resistant to primary breast cancer growth in addition to superior control of spontaneous tumor metastasis. CISH deletion favors NK cell accumulation to the primary tumor, optimizes NK cell killing properties and decreases TIGIT immune checkpoint receptor expression, limiting NK cell exhaustion. Finally, we argue that specifically enhancing NK cell function is sufficient to boost anti-tumor response to both primary and secondary tumor models. Using CRISPRi, we then targeted *CISH* in human NK-92 or primary NK cells. According to the results in our mouse model, CISH deletion favors NCR signaling and anti-tumor functions in human NK cells. Our results validate CISH as an emerging therapeutic target to enhance NK cell immunotherapy.

## INTRODUCTION

Immunotherapy strategies aim to mobilize the patient’s immune defenses against his tumor cells,targeting in particular cytotoxic effectors (CD8^+^ T cells, γδ T cells and natural killer (NK) cells). Antibody-based approaches aim at blocking inhibitory receptors (immune checkpoint inhibitor, ICI) on the surface of cytotoxic cells (CTLA-4, PD-1). These new therapies are mainly targeting CD8^+^ T cells and have a significant impact on the prognosis and survival of cancer patients. However, many patients are refractory to these treatments. The success and limitations of current immunotherapies have pushed research towards the development of alternative approaches and the possibility to manipulate other cytotoxic immune cells such as NK cells.

NK cells receive a renewed interest in immunotherapy because they do not rely on antigen specificity like CD8^+^ T cells, hence displaying a broader reactivity to tumors^1, 2, 3^. The distinction between a healthy cell (to be spared) and a malignant cell (undesirable) is possible thanks to a panel of inhibitory and activating surface receptors capable of integrating danger signals. NK cells also play a critical role in helping innate and adaptive immune responses via cellular cross-talk in various disease settings ^4^. Finally, in an adoptive transfer therapy setting, NK cells have greater off-the-shelf utility and are safer. Indeed, these cells are less persistent reducing the risk of autoimmunity and side effects such as cytokine storms (CRS, cytokine-release syndrome) that may result from an over activation of T cells. Therefore, therapeutic approaches that trigger and/or reconstitute NK cell function and proliferation are crucial in tumor immunotherapy. In addition, NK cells engineered with chimeric antigen receptor (CAR), targeting tumor antigens, are highly promising^5^.

In numerous preclinical studies, NK cells have shown an ability to limit the metastatic spread of experimental and spontaneous tumors^6, 7^. In the clinical setting, NK cell activity has been inversely correlated with cancer incidence ^8^, and several studies show that NK cell infiltration in lung, gastric, neuroblastoma and colorectal cancers is associated with better patient outcomes ^9, 10^.

Targeting inhibitory intracellular proteins, such as CISH, a member of the SOCS (Suppressor of Cytokine Signaling) family, can be effective in enhancing CD8^+^ T cell tumor immunity in preclinical models ^11, 12^. These studies led to a clinical trial protocol NCT04426669. CISH has been also shown to be a critical immune checkpoint in NK cells ^13^ and we recently contributed to show the importance of CISH in NK cell homeostasis using a *Cish* KO germline mouse model ^14^. However, we still do not fully appreciate mechanistically how CISH regulates NK anti-tumor functions. These are essential requirements to comprehensively uncover the safety of genetically or pharmacologically targeting of an intracellular protein in NK cell immunotherapy.

To further understand how CISH regulates NK cell activity, we developed and investigated a new conditional mouse model that depletes CISH specifically in NK cells (*Cish^fl/fl^Ncr1^iCre^*). In conditional *Cish*-deficient mice (*Cish^fl/fl^Ncr1^iCre^*), we found no change in homeostatic NK cell numbers, no alterations in maturation or cycling. Global gene expression of *Cish^fl/fl^Ncr1^iCre^* NK cells compared to *Cish^+/+^Ncr1^iCre^* NK cells revealed upregulation of pathways and genes associated with NK cell cycling and activation. Indeed, *Cish^fl/fl^Ncr1^iCre^* NK cells displayed a lowered activation threshold upon IL-15 *ex vivo* stimulation. We showed that CISH expression is not only triggered by cytokine stimulation but also upon Natural Cytotoxicity Receptor (NCR) stimulations that are essential for tumor cell recognition. Upon NCR triggering, we showed enhanced functions and signaling in *Cish*-depleted NK cells. Morevover, *Cish^fl/fl^Ncr1^iCre^* mice were more resistant to experimental tumor metastasis and primary tumor growth. The latter result showed for the first time, that NK-cell specific deletion of CISH in mature NK cells can dramatically enhance their anti-tumor function in primary tumors. Interestingly, we showed that *Cish*-depleted NK cells are more frequent in solid tumor and expressed less of the inhibiting receptor TIGIT *in vivo* and *ex vivo,* reducing NK cell exhaustion. We conclude that CISH targeting improves NK cell proliferation, functions, activation of several signaling pathways (cytokine receptors, but also NCRs) and limits NK cell exhaustion. This study represents a crucial step in the mechanistic understanding and safety of *Cish* targeting to unleash NK cell anti-tumor function in solid tumors. Finally, using a CRISPRi method, we were able to efficiently target *CISH* in NK-92 and primary NK cells improving their anti-tumor properties.

## RESULTS

### Deletion of CISH in mature NK cells does not impact maturation

To develop a conditional *Cish*-deficient mouse, C57BL/6 germplasm (sperm) were obtained with a Knockout First allele with conditional potential (Cish^tm1a(KOMP)Wtsi^). The EPD0165_3_H07 sperm clone was injected into C57BL/6 female mouse. Mice containing the germline-transmitted Cish^tm1a^ allele were confirmed by PCR and bred to a FLP deleter strain to remove the *lacZ* and *neo^r^* genes flanked by FLT recognition target sites to generate the floxed *Cish* allele (*Cish*^tm1cCiphe^ or *Cish*^fl/fl^). *Cish*^fl/fl^ mice were then bred with the Ncr1^iCre^ transgenic mice to induce deletion of *Cish* in NK cells (Supplemental Figure 1A). Efficient deletion of CISH was first confirmed by western blot and showed no evidence of CISH in purified and expanded splenic NK cells stimulated with IL-2 or IL-15 (supplemental Figure 1B). Similar to germline *Cish*-deleted mice, *Cish^fl/fl^Ncr1^iCre^* mice were healthy, fertile and contained equivalent frequencies and numbers of NK cells in bone marrow (BM) and spleen (Supplemental Figure 1C-E).

To determine whether *Cish*-deletion in NK cells perturbs their biology, we assessed the phenotype of NK cells that were deleted of *Cish* from the NKp46^+^ stage onwards. We did not detect NCR expression differences nor CD122 (IL-15Rβ) and DNAM1 (suppl. Fig 2 A-C). The maturation of *Cish^fl/fl^Ncr1^iCre^* NK cells is normal compared with *Cish^+/+^Ncr1^iCre^* NK cells showing equivalent frequencies of immature, M1 and M2 NK cell populations according to CD11b and KLRG1 (Figure 1A-B) or CD27 and CD11b expression in spleen and BM (Supplemental Figure 2D-E). Altogether, we did not notice any phenotypical differences in *Cish^fl/fl^Ncr1^iCre^* NK cells. This result confirms that targeting CISH is not triggering any adverse effects or altering NK cell subset phenotype at steady state. We next compared the gene expression profiles of *Cish^+/+^Ncr1^iCre^* versus *Cish^fl/fl^Ncr1^iCre^* NK cells.

**Figure 1.**
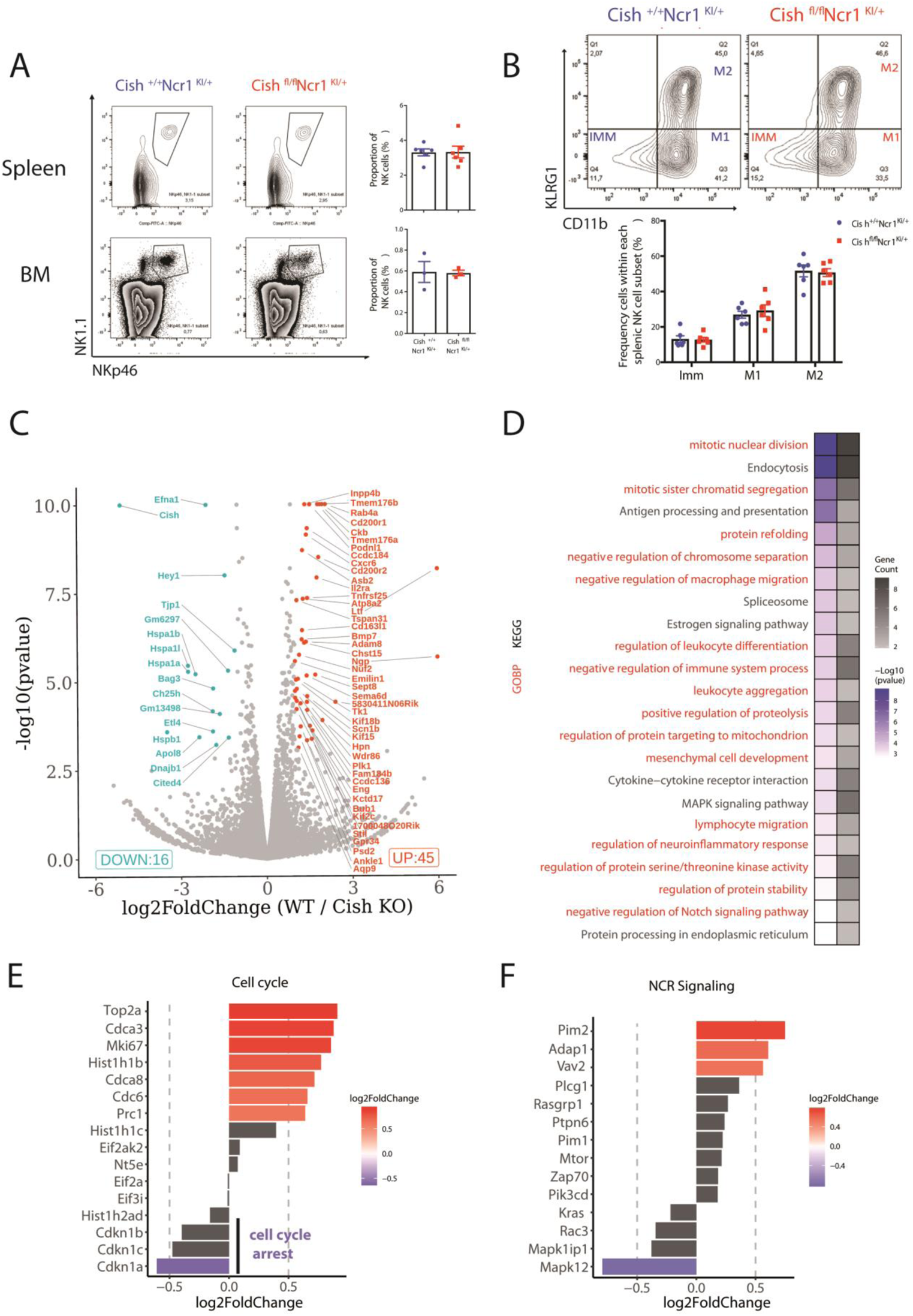
Deletion of Cish gene in mature NK cells does not impact NK cell maturation but depletion reveals upregulation of pathways and genes associated with NK cell cycling and activation. (**A-B**) BM and/or splenic NK cells from *Cish^+/+^Ncr1^Ki/+^* and *Cish ^fl/fl^Ncr1^Ki/+^* mice were phenotypically analysed by flow cytometry. (A) Frequency of NK cells in spleen and bone marrow (CD3^−^, CD19^−^, NK1.1^+^, NKp46^+^), representative FACS plots and histograms are shown. (B) Frequency of Imm, M1 and M2 cells were measured within NK cell populations from the spleen, representative FACS plots and histograms are shown. (**C-F**) Gene expression profiles of FACS sorted NK cells (NK1.1^+^, NKp46^+^, CD3^−^) were generated using RNA-sequencing. (C) A volcano plot of the top 60 significant (*p*<0.05) differentially expressed genes with a fold change above +1 or below −1 was generated. y-axis is the negative log-*p*-value and x-axis is the log-fold-change of the corresponding gene in *Cish^−/−^* vs *Cish^+/+^* comparison. Gene expression: blue = downregulated, red = upregulated. (D) Heatmap showing results of gene ontology analysis for genes in *Cish^−/−^* vs *Cish^+/+^* comparison (From KEGG pathway analysis in black or GOBP in red), log-*p*-value are indicated in blue color gradient and gene count in black color gradient. An upregulated list of genes associated with cell cycle (E) and Natural Cytotoxicity Receptor pathways (F) (from KEGG pathway analysis) was generated with a *p* value cut-off of <0.05. x-axis is the log(FC) of each gene and *p* values are indicated by colour.

### CISH depletion reveals upregulation of pathways and genes associated with NK cell-cycling and activation

Naïve *Cish^+/+^Ncr1^iCre^* or *Cish^fl/fl^Ncr1^iCre^* cells from spleen were purified and analyzed by RNAseq. After *Cish* deletion, more than 200 genes were differentially expressed (+/- 20% expression, adjusted p-value ≤ 5%). Among them, about 60 genes have a modulated expression of a fold increase greater than 2 (Fig.1C). An analysis by KEGG and GO enrichment targeting signaling pathways such as cell cycle progression, cytokine signaling or NK activating receptors, allowed us to assess the influence of the absence of *Cish* on these biological processes and to determine the putative markers responsible for these perturbations (Fig.1 D-F). Overall *Cish^fl/fl^Ncr1^iCre^* cells appeared to display a more-activated phenotype than their *Cish^+/+^Ncr1^iCre^* counterparts. Subsequently, we decided to evaluate cellular functions of *Cish*-deficient NK cells.

### CISH depletion favors IL-15 cytokine signaling pathway decreasing NK activation threshold

Following IL-15 stimulation, *Cish*-depleted NK cells exhibited an increased proliferation compared to *Cish^+/+^Ncr1^iCre^* cells (Fig. 2A-B). Interestingly, *Cish^fl/fl^Ncr1^iCre^* cells proliferate efficiently at IL-15 doses as low as 10ng/ml, while *Cish^+/+^Ncr1^iCre^* cells are almost not proliferating. We detected no proliferation of *Cish^+/+^Ncr1^iCre^* and *Cish^fl/fl^Ncr1^iCre^* cells at 1ng/ml IL-15 doses, suggesting that *Cish^fl/fl^Ncr1^iCre^* cells can probably proliferate efficiently at IL-15 doses between 1 and 10 ng/ml. *Cish^fl/fl^Ncr1^iCre^* cells showed max proliferation rate at 30ng/ml of IL-15, whereas *Cish^+/+^Ncr1^iCre^* cells needed at least 50ng/ml. These results show that *Cish^fl/fl^Ncr1^iCre^* NK cells are more sensitive to IL-15 and their activation threshold is greatly diminished in absence of *Cish*. We next decided to follow CD122 (IL-15Rβ) receptor expression during IL-15 activation. CD122 expression remained constant during 6 days in a *Cish^fl/fl^Ncr1^iCre^* context while it decreased in *Cish^+/+^Ncr1^iCre^* at day 3 and 6 (Fig. 2C, D). This result suggests a role for CISH in the recycling and degradation of CD122 receptor. We also observed that IL-15-stimulated *Cish*-deficient NK cells showed improved IFN-γ and CD107a expression (Fig. 2E, F). This result correlates with previous results obtained in germline deficient mice ^13^. In the absence of CISH, following cytokine stimulation, there is no negative feedback and NK cell functions are amplified. The role of CISH under other NK signaling pathways has not been explored.

**Figure 2:**
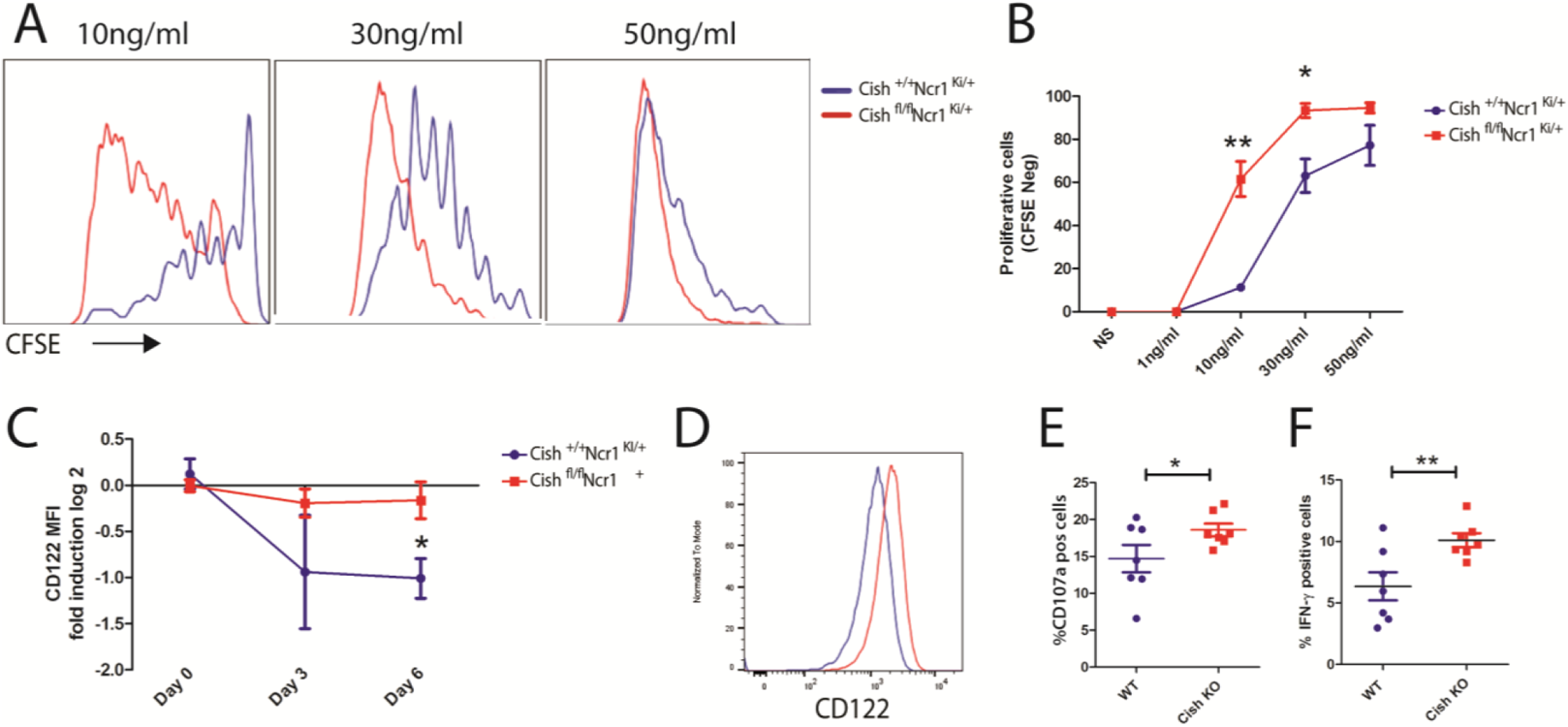
Cis depletion favors IL-15 cytokine signaling pathway decreasing activation threshold. **(A-B)** Flow cytometric analysis of *Cish ^+/+^Ncr1^Ki/+^* and *Cish ^fl/fl^Ncr1^Ki/+^* purified NK cells labeled with carboxyfluorescein succinimidyl ester (CFSE) and cultured for 6 d in IL-15 (10–50 ng/ml). (A) Representative FACS plot is showed. (B) Representative curve of 3 different experiments are shown. **(C-D)** Flow cytometry was used to quantify the geometric mean of CD122 expression at different time points on *Cish ^+/+^Ncr1^Ki/+^* and *Cish ^fl/fl^Ncr1^Ki/+^* NK cells. (C) Representative curve of 3 different experiments are shown. (D) Representative FACS plot is showed. **(E-F)** Histograms of 6 different experiments Flow cytometric analysis of IFN-γ production (E) and CD107a (LAMP-1) (F) expression *Cish ^+/+^Ncr1^Ki/+^* and *Cish ^fl/fl^Ncr1^Ki/+^* NK cells from splenocytes cultured for 4 hrs in IL-15 (50ng/ml). \**p<*0.05, \*\**p<*0.01 (Student’s *t*-test).

### CISH is expressed upon Natural cytotoxicity receptors stimulation but mildly affected naïve NK cell functions and signaling

We next decided to determine CISH expression upon cytokine stimulation. As expected, CISH expression is induced after 4hrs of IL-2 or IL-15 stimulation (Fig. 3A). Interestingly, CISH expression is almost undetectable without stimulation. NK cells express a variety of receptors that allow them to detect target cells (endangered, infected or tumor cells) while sparing normal cells ^15^. This recognition is performed via a multitude of activating receptors such as the natural cytotoxicity receptors (NCRs) (NKp30, NKp44, NKp46), NKG2D, 2B4, DNAM-1 or the Fc fragment receptor for immunoglobulins, CD16 (FcRIIA) ^3^.

**Figure 3:**
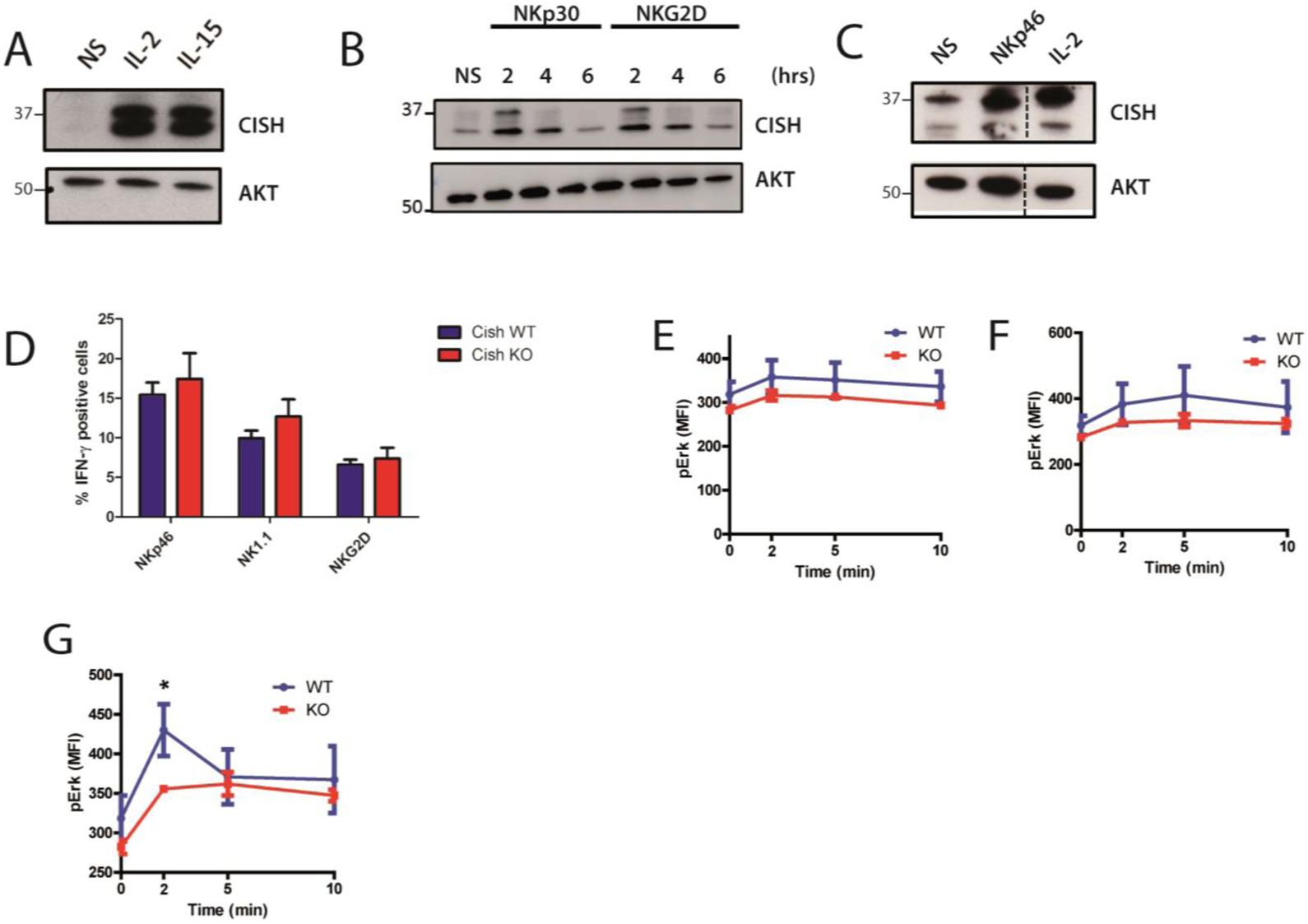
CISH is expressed upon natural cytotoxicity receptors stimulation but mildly affects naïve NK cell functions and signaling. **(A-B)** Human KHYG-1 NK cells were rested O/N, then stimulated for 4hrs with IL-2 (3000 UI/ml) and IL-15 (50ng/ml) (A), or coated anti-NKp30 or anti-NKG2D antibody (2μg/ml) for the indicated time (B). Cell lysates were analysed by immunoblotting for CISH or AKT (loading control). **(C)** Spleen purified NK cells were stimulated for 4 hrs with IL-2 (3000 UI/ml) or coated anti-NKp46 antibody. Cell lysates were analysed by immunoblotting for CIS or Akt (loading control). **(D-G)** Total splenic cells were harvested from 6-8 week-old *Cish^+/+^* and *Cish^−/−^* mice and depleted of red blood cells. (D-E) Cells were cross-linked with anti-NK1.1, anti-NKp46 and anti-NKG2D coated antibodies for 4 hrs in the absence of IL-15. Flow cytometric analysis of NK cells and their IFN-γ production (D) was assessed after 4 hours. (E-G) Cells were cross-linked with anti-NK1.1, anti-NKp46 and anti-NKG2D coated antibodies for 2, 5 and 10 minutes. Flow cytometric analysis of ERK1/2 phosphorylation status in NK cells was assessed.

The signaling pathways induced by the involvement of TCR (T Cell Receptor) and NCRs (Natural Cytotoxicity Receptors) are quite similar^16^. We hypothesized that CISH could be involved in the signal encoded by NCR engagement as suggested by our RNA-seq analysis (Fig.1F). We showed that CISH can be induced following stimulation of NKG2D, NKp30 or NKp46 activating receptors, respectively in a human NK lineage (NK-92) and naive mouse NK cells (Fig. 3 B-C). These receptors and the integrity of their signaling pathways are essential for the recognition and subsequent elimination of a tumor cell.

The enrichment of genes downstream of activating receptors such as NK1.1, NKp46 and NKG2D, suggests a more activated state of *Cish*-deficient NK cells *in vivo*. We next decided to test NK cell cytokine expression upon NCR activation. Prior to this, we confirmed that there are no differences in activating receptor expression on the surface of *Cish^+/+^Ncr1^iCre^* versus *Cish^fl/fl^Ncr1^iCre^* NK cells (suppl. Fig 2C). We stimulated naïve *Cish^+/+^Ncr1^iCre^* or *Cish^fl/fl^Ncr1^iCre^* NK cells with plate bound NKp46, NK1.1 or NKG2D antibodies (Fig. 3D). We detected no statistical difference in IFN-γ expression, although *Cish^fl/fl^Ncr1^iCre^* cells tend to express more of IFN-γ. The same results were obtained with stronger stimulations such as PMA (phorbol myristate acetate) and ionomycin (suppl. Fig 3A). We then decided to explore naïve *Cish^+/+^Ncr1^iCre^* and *Cish^fl/fl^Ncr1^iCre^* NK cells signaling pathway after short NKp46, NK1.1 or NKG2D cross-linking stimulations (Fig. 3E-G). Phosphoflow analyses were performed to detect ERK phosphorylation status,as this signaling pathway is induced upon NCR engagement. A slight increase of ERK phosphorylation is detected for all stimulations, but we detected no statistical difference in phospho-ERK increase between stimulated *Cish^+/+^Ncr1^iCre^* and *Cish^fl/fl^Ncr1^iCre^* naïve NK cells. We did not detect any difference as well when stimulating with PMA and Ionomycin (suppl. Fig3.B). Altogether, no major differences are detected in naïve NK cells upon NCR triggering, which might be due to the fact that CISH is weakly expressed at steady state. As primed NK cells are more susceptible to express CISH, similar experiments were performed in these cells.

### CISH deletion favors *ex vivo* expanded NK cell signalling, proliferation and functions

Activation and amplification of splenic murine NK cells (LAKs; Lymphokine-activated killer cells) have been previously described^17^. Briefly, total splenocytes are cultured in high IL-2 concentration (1000 UI/ml). At day 6, we had more total LAK cells in *Cish^fl/fl^Ncr1^iCre^* compared to *Cish^+/+^Ncr1^iCre^* genotype (Fig. 4A). More interestingly, we were able to produce twice the total absolute number of NK cells in *Cish^fl/fl^Ncr1^iCre^* condition compared to *Cish^+/+^Ncr1^iCre^* (Fig 4B). At day 6 post-IL-2 stimulation, only three kinds of cell remain in the culture:, NK cells (NK1.1+, CD3−), NKT cells (NK1.1+, CD3+) and T cells (NK1.1-, CD3+). We observed more production of NK cells and NKT cells in *Cish^fl/fl^Ncr1^iCre^* splenocytes cultures and as a counterpart a dramatic decrease in T cell proportion (Fig 4C). NK cells and NKT cells both express NKp46 receptor and are thus Cish-deficient in *Cish^fl/fl^Ncr1^iCre^* mice. Since we showed that Cish-deficient NK cells are more sensitive to cytokine stimulations, this result is in accordance with our previous observations.

**Figure 4:**
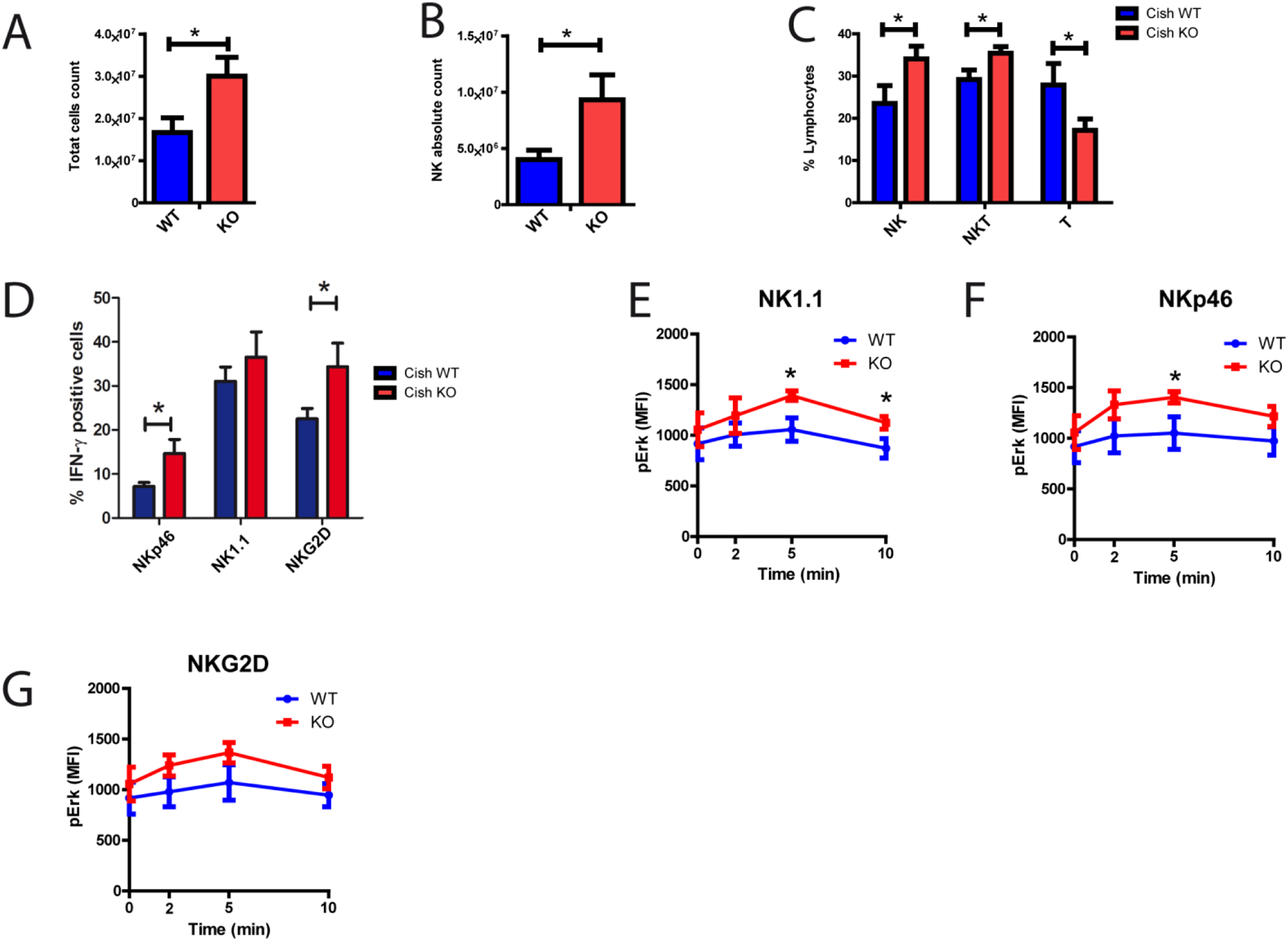
Cis deletion favors *ex vivo* expanded NK cells proliferation, functions and signaling. **(A-G)** Total splenocytes were expanded in IL-2 (1000UI/ml). (A) Histograms showing absolute numbers of cells that have proliferated at day 6. (B) NK cells gated by FACS (NK1.1+, CD3–) absolute count. (C) FACS gated histogram percentage of NK cells (NK1.1+, CD3−), NKT cells (NK1.1+, CD3+) and T cells (NK1.1−, CD3+). (D-F) Upon an overnight IL-2-starvation, cells were stimulated with anti-NK1.1, anti-NKp46 and anti-NKG2D coated antibodies for 4 hrs. Flow cytometric analysis of NK cells and their IFN-γ production (D) was assessed after 4 hrs. (E-G) Cells were cross-linked with anti-NK1.1, anti-NKp46 and anti-NKG2D coated antibodies for 2, 5 and 10 minutes. Flow cytometric analysis of ERK1/2 phosphorylation in NK cells was assessed. \**p<*0.05, \*\**p<*0.01 (Student’s *t*-test).

We next decided to test whether *Cish^fl/fl^Ncr1^iCre^* NK cells were more sensitive to activating receptor stimulation [NKp46, NK1.1, NKG2D] than their *Cish^+/+^Ncr1^iCre^* counterparts.We did not detect any receptor expression differences except for NKG2D that was surprinsingly less expressed in *Cish^fl/fl^Ncr1^iCre^* NK cells compared to *Cish^+/+^Ncr1^iCre^* NK cells, in contrast to CD122 that was more expressed as previously described (suppl. Fig. 3C-F). *Cish^fl/fl^Ncr1^iCre^* and *Cish^+/+^Ncr1^iCre^* NK cells from LAK cultures were stimulated with plate bound NKp46, NK1.1 or NKG2D antibodies. Primed *Cish^fl/fl^Ncr1^iCre^* NK cells showed more expression of IFN-γ especially upon NKp46 and NKG2D stimulations (Fig. 4D). We did not detect any difference as well when stimulating with PMA and Ionomycin (suppl. Fig. 3G). We then decided to investigate signaling in primed-NK cells using phosphoflow experiments, upon short NKp46, NK1.1 or NKG2D cross-linking stimulations (Fig.4 E,F,G). Here again, we were able to see a clear increase in ERK phosphorylation in *Cish^fl/fl^Ncr1^iCre^* NK compared to *Cish^+/+^Ncr1^iCre^* after NCRs stimulation, that was also detected using strong stimulation such as PMA and Ionomycin (suppl. Fig. 3H). These observations suggest that the loss of CISH lowers the activation thresholds of these activating receptors by acting on their signaling. This could lead to an increased anti-tumor potential of NK cells in the absence of *Cish*.

### Specific deletion of CISH in NK cells enhances immunity to metastasis

Given the increased activation and proliferation observed *in vitro*, we next asked whether deletion of CISH in NK cells could confer enhanced tumor immunity *in vivo*. We challenged *Cish^fl/fl^Ncr1^iCre^* and *Cish^+/+^Ncr1^iCre^* mice with two syngeneic tumor cell lines, known to be controlled by and stimulate NK cell activity. Intravenous (i.v.) administration of B16F10 melanoma cells showed a decrease in metastatic nodules in the lung compared to *Ncr1^iCre^* controls (Fig. 5A). Similarly, when E0771-GFP/Luc breast cancer cells were administered i.v. to *Cish^fl/fl^Ncr1^iCre^* and *Cish^+/+^Ncr1^iCre^* mice, tumor burden was significantly reduced at day 7 and 14 in *Cish^fl/fl^Ncr1^iCre^* mice compared to *Cish^+/+^Ncr1^iCre^* mice (Fig. 5B-C). The absence of CISH in NK cells also had an effect on the metastasis of E0771-GFP/Luc breast cancer cells to other organs. Indeed, metastases in the lungs of *Cish^fl/fl^Ncr1^iCre^* mice were significantly decreased compared to *Cish^+/+^Ncr1^iCre^* mice (Fig. 5D). E0771 metastasis to the liver was also measured, with *Cish^fl/fl^Ncr1^iCre^* mice exhibiting significantly reduced metastatic burden (Fig. 5E). We next decided to test efficiency of CISH deleted NK cells in a solid tumor model experiment.

**Figure 5:**
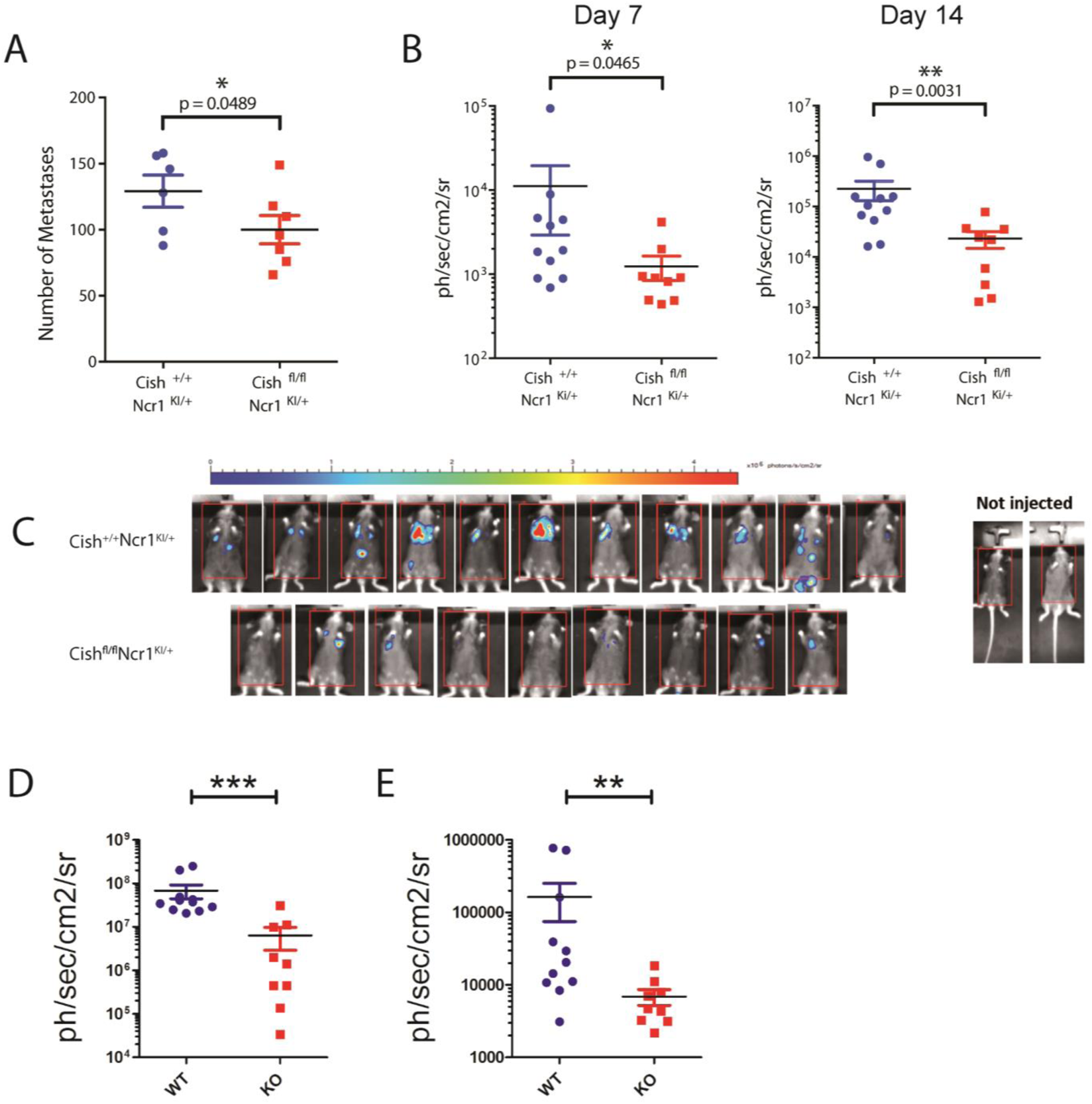
Specific deletion of CIS in NK cells enhances immunity to metastasis. **(A)** 3×10^5^ B16F10 melanoma cells were injected i.v. into *Cish ^+/+^Ncr1^Ki/+^* and *Cish ^fl/fl^Ncr1^Ki/+^* mice and 14 days later lung metastases were enumerated. (**B-C**) 5×10^5^ EO771-GFP^+^ Luciferase^+^ breast cancer cells were injected i.v. into *Cish ^+/+^Ncr1^Ki/+^* and *Cish ^fl/fl^Ncr1^Ki/+^* mice and metastatic burden was quantified by luminescence at day 7 (left panel) and 14 (right panel). (C) Representative luminescence pictures are showed. (**D, E**) 5×10^5^ EO771-GFP^+^ Luciferase^+^ breast cancer cells were injected i.v. into *Cish ^+/+^Ncr1^Ki/+^* and *Cish^fl/fl^Ncr1^Ki/+^* mice and spontaneous metastatic burden was quantified by luminescence at day 14 in the lung (D) and liver (E). \**p*≤0.05 \*\**p*≤0.01 \*\*\**p*≤0.001 (unpaired Student’s *t*-test).

### Specific deletion of CISH in mature NK cells enhances immunity to primary tumors

NK cells are best defined for their anti-metastatic function. Remarkably, we found that the growth of orthotopic E0771 tumor implanted in the mammary fat pad was also significantly reduced in *Cish^fl/fl^Ncr1^iCre^* mice compared to *Cish^fl/fl^Ncr1^iCre^* mice (Fig. 6A-B). This coincided with reduced spontaneous metastasis to the lungs (Fig. 6C-D). We also found more absolute number of infiltrated NK cells in the tumor (Fig. 6E).

**Figure 6:**
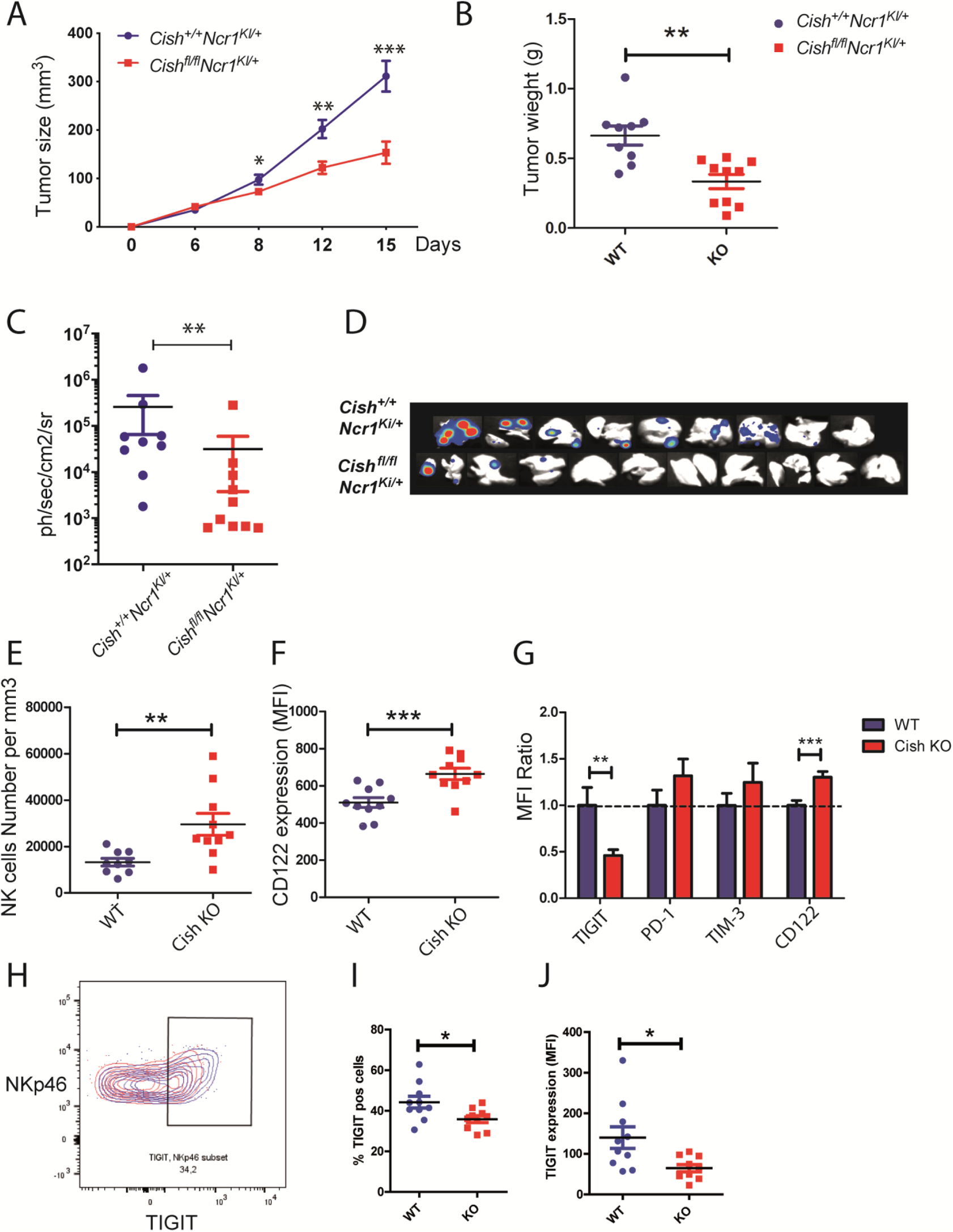
Specific deletion of CIS in mature NK cells enhances immunity to primary tumors. (**A-J**) Orthotopic injection of EO771-GFP^+^ Luciferase^+^ breast cancer cells in the mammary fat pad of *Cish ^+/+^Ncr1^Ki/+^* and *Cish^fl/fl^Ncr1^Ki/+^* mice. (**A**) Measure of tumor growth in mm^3^ at indicated timepoints. **(B)** Measure of tumor weight at day 15. **(C-D**) At day 15, lungs from tumour bearing *Cish ^+/+^Ncr1^Ki/+^* and *Cish^fl/fl^Ncr1^Ki/+^* mice were analysed and metastatic burden quantified by luminescence. **(D)** Pictures of lung luminescence at day 15. **(E)** Numbers of infiltrated NK cells (NK1.1^+^, CD3^−^) /mm^3^ of tumors. **(F)** Flow cytometry was used to quantify the mean fluorescence of CD122 expression on Infiltrated NK cells. **(G)** Flow cytometry was used to quantify the mean fluorescence ratio to WT infiltrated NK cells MFI of TIGIT, PD-1, TIM-3 and CD122. **(H-J)** TIGIT expression analysis by flow cytometry. (H) Representative plot comparing *Cish^+/+^Ncr1^Ki/+^* and *Cish^fl/fl^Ncr1^Ki/+^* TIGIT profile. **(I)** Percentage of TIGIT positive cells and **(J)** Mean fluorescence intensity of TIGIT. Mean ± s.e.m. of *n*≥9 biological replicates;\**p<*0.05, \*\**p<*0.01, \*\*\**p<*0.01 (Student’s *t*-test).

We also checked for activation of these cells, and observed no difference in CD69, CD25 expression, but also no difference in NK cells subtypes (KLRG1, CD11b) present in the tumor when comparing CISH KO NK cells with WT Cells (Suppl. Fig. 4A-D). However, we found a higher expression of CD122 (IL15-Rβ) receptor at the surface of CISH KO NK cells compared to WT cells (Fig. 6F), in accordance with the *in vitro* observations (Fig. 2D).

One of the main causes of relapse in immunotherapy treatment is the exhaustion of cytotoxic cells due to inhibiting receptor expression. We thus tested for expression of the PD-1, TIM-3 and TIGIT receptors that are all suggested to be a part of NK cell exhaustion ^18^ (Fig. 6G). Interestingly, only TIGIT expression was strongly decreased in infiltrated CISH KO NK cells compared to WT NK cells (Fig. 6H-J). We then wanted to see if this observation was confirmed in infiltrated NK cells in human breast cancer patients. For this, we used previously published Database of Single cells mapping of immune cells infiltrated in human breast tumors^19^. Using loupe cell browser software, infiltrated NK cells (*CD3*-, *NCAM1+*) were gated then *TIGIT* gene expression was compared in *CISH^+^* NK cells versus *CISH^−^* cells. Interestingly, *TIGIT* is one of the top ten up-regulated gene in presence of *CISH* (suppl. Fig. 4E) and that *TIGIT* gene expression was greatly reduced when *CISH* gene is absent (suppl. Fig. 4F). This latter result suggests that *TIGIT* and *CISH* are regulated similarly in NK cells.

These data highlight a previously unappreciated role for NK cells in controlling the growth of primary tumors in addition to metastatic cancer cells. We are also showing for the first time, that this efficiency may be the results of several factors especially increased sensitivity to cytokines present in the TME, the tumor cell recognition and less exhaustion. These data strengthen the rationale for the targeting of CISH in human NK cells to improve cancer immunotherapy.

### Targeting *CISH* genetically in human NK cells using CRISPR(i)-Cas9 technology improves their function

The use of human NK cells in adoptive transfer therapy has showed promising results^20^. We decided to target human NK-92 cell line using the cutting edge tool CRISPR(i)-dCas9 to target *CISH* ^21^. This method represses gene-targeted promoter and reduce drastically potent “off-target” effect and allows a potential reversible method. NK-92 cells were transduced with a KRAB-dCAS9 construct co-expressing mCherry and a control or *CISH* sgRNA construct co-expressing GFP (Fig. 7A). We used the classical lentivirus envelop VSV-G to produce virus particles and subsequently transduce NK-92 cells. mCherry and GFP double positive cells were selected by cell sorting. CISH expression is reduced in sgCISH transduced cells compared to control NK-92 transduced cells (Fig. 7B). We then investigated the reactivity of transduced NK cells. In a co-culture assay, sgCISH NK-92 cells led to an increased apoptosis of K562 target cells compared to control transduced NK-92 cells (Fig. 7C). Similarly, sgCISH NK-92 cells show higher level of degranulation (CD107a+) and production of TNF-α compared to control transduced NK-92 cells (Fig 7D). We showed previously that CISH is expressed upon NKp30 stimulation in NK-92 cells. We thus hypothesized that CISH may regulate NKp30 signaling. Control and sgCISH transduced cells were stimulated with cross-linked NKp30 antibody during indicated time (Fig. 7E), WB analyses were performed showing an increase of ERK-1/2 and AKT phosphorylation in cells transduced with sgCISH compared to control. This difference is not due to NKp30 expression difference (suppl. Fig. 5A). We also checked for TIGIT expression in these genetically modified cells and observed less TIGIT expression when CISH is downregulated (Fig. 7F), in coherence with previous results observed in tumor infiltrated mouse NK cells (Fig. 6G-J).

**Figure 7:**
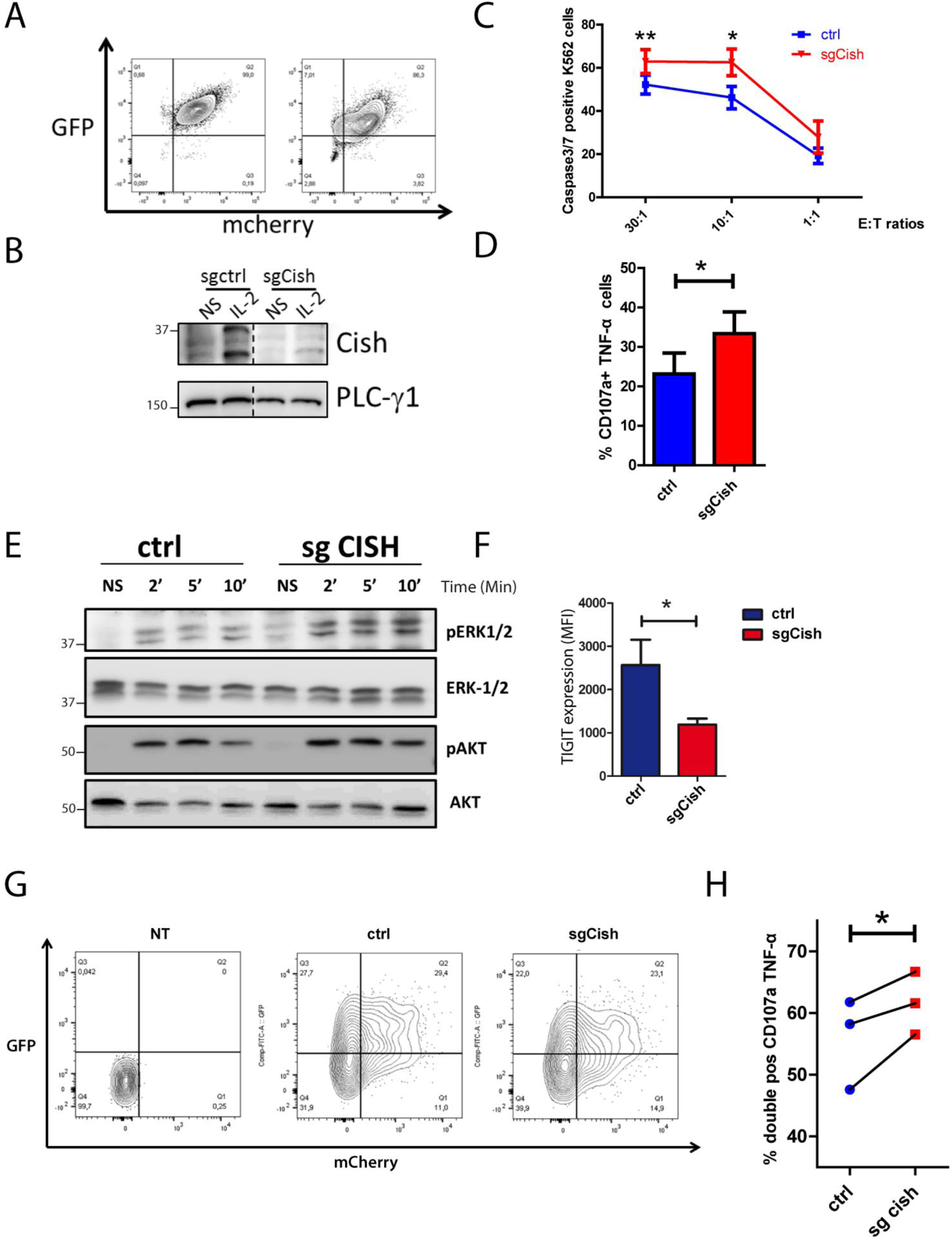
NK-92 / primary NK cells were transduced with a KRAB-dCAS9 construct co-expressing mCherry and a ctrl or Cish SgRNA construct co-expressing GFP. (**A**) Validation of constructs expression by flow cytometry in NK-92 cells. (**B**) Validation of CISH silencing by western blot, following IL-2 stimulation for 4hrs in NK-92 cells. (**C-D**) 4hrs co-culture of NK-92 WT or KO cells with K562 cancer cells. (**C**) cytotoxicity tests, K562 cells positive for caspase 3/7 marker are identified by flow cytometry. (**D**) Percentage of NK-92 cells double positive for CD107a and TNF-α identified by flow cytometry. (**E**) Upon an overnight IL-2 starvation, cells were stimulated with biotinylated NKp30 antibody (10 μg/ml) then cross-linked with streptavidin (20 μg/ml). Cell lysates were analysed by immunoblotting for p(T202/Y204)-ERK-1/2, ERK-1/2, p(S473)-AKT or AKT. (**F**) TIGIT expression was assessed by flow cytometry comparing NK-92 control (ctrl sg) and CISH-KO NK-92 (Sg CISH). (G) Validation of constructs expression by flow cytometry in primary NK cells 2 days after transduction. (H) 4hrs Co-culture assay of sgCISH or control transduced primary NK cells with K562 cancer cells at a ratio of 1:30 effector:target. Percentage of double positive cells for CD107a and TNF-α was identified by flow cytometry.

We next decided to use these validated construct in primary NK cells from blood of healthy volunteers. Development of immunotherapies targeting NK cells has been limited in part due to their resistance to traditional viral gene delivery systems. Indeed, in NK-92 experiments using classical lentiviral vectors pseudotyped with the Vesicular stomatitis virus G (VSV-G) envelope we achieved only low transduction efficiency. In order to overcome this poor transduction efficiency we decided to use the endogenous baboon retroviral envelope for pseudotyping of the lentiviral vectors that showed recently impressive results fo primary NK cells transduction ^22^. Using a MOI of 2, we were able to achieve around 50% of transduction efficiency for KRAB-dCAS9 construct co-expressing mCherry and ctrl or *CISH* sgRNA construct co-expressing GFP (Fig. 7G). We also achieved between 20 and 30 % of double positive cells. After sorting double positive cells, we performed a co-culture assay with K562 cells (Fig. 7H). We showed that primary NK cells from different donors transduced with sgCISH express more CD107a+ and TNF-α compared to control tranduced cells, this without affecting expression of surface receptors TIGIT, NKp30, NKG2D or CD122 (suppl. FIG 5 G-J). This latter result confirmed our results obtained with NK-92 cells.

Altogether, we showed the feasibility to target CISH in human NK cells using a technology combining the CRISPR(i)-dCas9 tool with a new lentiviral pseudotype allowing high level transduction efficiency in primary NKs. This method may be of great interest in the future to engineer primary NK cells. Indeed, we confirm that CISH is an important intracellular checkpoint inhibitor to target in clinic and describe in our study important tools that may be use to achieve this purpose.

## DISCUSSION

Many of the current treatments focus on targeting inhibitory cell surface receptors to improve cytotoxic lymphocyte anti-tumor function. Here, we suggest that targeting intracellular “brakes” such as CISH may also be of beneficial interest. Considering the off-target effects of many current immunotherapy treatments, we are mindful of examining the basic biology of targeting intracellular checkpoints alongside the potent anti-tumor responses prior to clinical translation of this strategy. This study explores in detail the effects of depleting CISH in NK cells specifically (*Cish^fl/fl^Ncr1^iCre^)*.

*Cish*-deficient cells are terminally mature and we did not observe any differences in NK cell development in NK cells isolated from *Cish^fl/fl^Ncr1^iCre^* mice compared to WT NK cells. This is contrasting with the maturation defect and DNAM-1 over expression that was found on germline CISH-deleted NK cells ^14^. We believe that CISH regulates these receptors expression processes at an earlier NK cell stage prior to NKp46 expression. Indeed, targeting CISH using CRISPR-Cas9 technology in iPSCs-derived NK cells lead also to a delay in NK cell differentiation ^23^. This suggests that targeting CISH in mature NK cells may be a safe strategy in the clinic preventing potential developmental defects. Our gene set enrichment analysis (GSEA) shows upregulation of signaling pathways essential for NK cell activation and cycling. This observation indicates that in absence of CISH, NK cells are “programmed” to proliferate actively and to be activated. This GSEA allowed us to suggest that CISH may also be a part of other signaling pathways and not only cytokines pathways. Indeed, like we showed in T cell, NK cells increased activation is the result of several signaling pathways i.e. cytokines but also NCR as we show in this study. CISH belong to suppressor of cytokines (SOCS) family and has been first described as a negative regulator of cytokine signaling. We showed in T cells previously and here in NK cells that CISH get a more complex role in cell signaling pathways, we thus believe that probably the other SOCS proteins may have broader effects on signaling as well. In naïve NK cells, we only saw a mild effect of CISH absence functionally. We believe this is due to the fact that CISH is expressed at a very low level at steady state, and will be expressed upon cytokine and or NCR stimulations. This is probably why we had a greater activation of our CISH KO NK cells compared to WT cells, especially after NCR activations. Indeed, we just stimulated these receptors one by one, but *in vivo* NK cell responses to infectious or tumor cells depend on the balance between several activating and inhibitory signals. Thus, the increased NK cells killing properties observed in absence of CISH is probably the addition of several over activated signaling pathways. NK cells become more sensitive to their environment due to the decrease in their activation threshold.

It was previously shown that CISH is a potent checkpoint in NK cell-mediated anti-metastatic effect using a CISH germline deficient mouse, *Cish^−/−^*^13^. However, CISH deletion in DC^24^, macrophages ^25^ and T cells^12, 26^ also favoured the activation of these immune cells. Here, we definitively show that the absence of CISH in NK cells is sufficient to decrease metastasis formation, by using a conditional CISH KO *Cish^fl/fl^Ncr1^iCre^* mouse model. The reduction of tumor metastases in this model emphasizes the importance of NK cells in the anti-tumor response and rules out the role of other immune cells deficient in CISH that may have influenced tumor regression in the germline model.

Previously, an orthotopic injection of E0771 breast cancer cells into germline *Cish^−/−^* mice resulted in decreased tumour burden and metastasis to other organs ^13^. However, there is limited evidence to suggest that the decrease in tumour size is due to increased NK cell function alone, or the orchestration of multiple immune cells lacking CISH (such as CD8+ T cells). Strikingly, when *Cish^fl/fl^Ncr1^iCre^* mice were orthotopically implanted with E0771 breast cancer cells, CISH deletion in NK cells alone was enough to decrease tumor burden over time.

We observed more NK cells infiltrated in tumors in absence of CISH. The reason to that might be multifactorial. One reason to that may be that they are more sensitive to IL-15 and thus may proliferate more efficiently in response to the IL-15 that is present is the tumour microenvironment, since we clearly show here that infiltrated CISH KO NK cells express more CD122 (IL15-Rβ). We cannot rule out in this study, whether Cish KO NK cells are more efficiently recruited to the tumor site. Further studies are required to test for Cish KO NK cells homing. We also show in here that activated NK cells in absence of CISH produce more cytokines and degranulate more, especially in response to NCR receptors that are essential for tumor recognition and killing. This means that these cells are more eager to recognize and to kill directly their targets but also probably to influence other immunes cells due to their cytokine’s secretion such as DCs and CD8^+^ T cells. Unexpectedly, we also observed that Cish KO NK cells express less TIGIT that WT cells. So far, TIGIT has been the only one well defined checkpoint inhibitor receptor involved in NK cell exhaustion^27^. This latter result indicates that Cish KO NK cells are less exhausted than their WT counterparts. Further studies are required to understand in detail if TIGIT and CISH are regulating each other expressions or if they are regulated by the same signaling pathways, as it has been suggested for example for IL-15 ^28^. Thus, targeting CISH in NK cells represents a promising strategy in immunotherapy not only for anti-metastatic indications but also in primary tumor progression.

NK cell immunotherapy holds great promise as an off-the-shelf cell therapy but their full potential has not been reached yet. Here we propose to target these cells with the elegant CRISPR(i)-dCas9 tool and show its feasibility in primary NK cells using the appropriate endogenous baboon retroviral envelope. Other protocols have been proposed to target NK cells from blood using other viruses or electroporation methods^29, 30^. But all these protocols are using the regular Cas9 construct cutting irreversibly the targeted gene. Here we show for the first time that repressing CISH transcription using dCas9-KRAB construct is sufficient to improve primary NK cells functions. This method not only prevents classical off-target effects, but also can be used reversibly to avoid potent undesired side effects^21^. Our study represents an important proof of concept in order to use this tool but more developments are required to refine this method in the future.

In conclusion, targeting CISH improves NK cell *ex-vivo* proliferation, functions and signaling activation of several pathways such as cytokines and NCRs. *In vivo* CISH absence favors NK cell numbers to the tumor burden, otpimize their killing properties and limit NK cell exhaustion. We finally propose a new method to efficiently target gene in primary human NK cells.

Thus, in this study we confirmed and further characterized the inhibition of CISH as a method to unleash the NK cell anti-tumor response. We propose that releasing NK cell inhibition by targeting CISH is a safe strategy to improve NK cell anti-tumor properties and supports the clinical development of this approach.

## METHODS

### Mice

To generate Tm1c *Cish^fl/fl^* mice, sperm from *Cish*^tm1a(KOMP)WTsi^ Knock-out mouse project repository (KOMP, UC Davis)^31^ was injected into C57BL/6N host embryos at the Centre d’Immunophénomique (Ciphe) (Marseille, France). Homozygous *Cish*^tm1a(KOMP)WTsi^ mice were then crossed with FLP-FRT mice to generate Tm1c *Cish^fl/fl^* mice (B6-Cish^tm1cCiphe^). Then, Tm1c *Cish^fl/fl^* mice were crossed with *Ncr1*^iCre/+^ mice ^32^. Male and female mice were used between the ages of 6–12 weeks. Age and sex matched mice were used and cohort size was dictated by previous experience using these tumor models. Mice were bred and maintained under specific pathogen-free conditions at the Centre de Recherche en Cancérologie de Marseille (CRCM) animal facility. Animal experiments followed were performed in accordance with institutional committees and French and European guidelines for animal care.

### Genotyping

Tm1c *Cish^fl/fl^* genotyping was performed using the following PCR primers: *Cish* Tm1c Fwd, 5’-GAGGTCTCCCTGAGAACCCC-3’; *Cish* Tm1c Rev, Cis2, 5’-TTCCGCCACTGAGCCACATA-3’; with expected band sizes at 305 bp for WT alleles and 460 bp for floxed allele. *Ncr1^iCre/+^* genotyping was performed using the following PCR primers: iCRE Fwd, 5’-GGAACTGAAGGCAACTCCTG-3’; iCRE Fwd KI, 5’-GTCCATCCCTGAAATCATGC-3’; Rev WT:-5’ TTCCCGGCAACATAAAATAAA-3’; with expected bands sizes at 300 bp for WT allele and 376 bp for KI allele.

### NK cell expansion and Western blots

NK cells were purified from mouse spleens using Mouse NK cell isolation Kit according to manufacturer’s specifications (STEMCELL #19855). NK cells were then expanded for 6 days with hIL-15 50ng/ml (Miltenyi #130-095-765) in RPMI 20% FCS supplemented with L-glutamine (2 mM; Gibco), penicillin/streptomycin (100 μg/mL; Gibco), Sodium Pyruvate (1 mM; Gibco) and finally FACS sorted (NK1.1^+^, NKp46^+^, CD3^−^, CD19^−^) (BD Biosciences FACSAria III). After overnight starvation in the absence of IL-15, *Cish^+/+^Ncr1^iCre^* or *Cish^fl/fl^Ncr1^iCre^* NK cells were stimulated or not during 4 hours with IL-2 (Proleukin, Novartis pharma SAS) or IL-15. Cells were lysed at 4 °C for 10 min in 1% NP-40 lysis buffer (50 mM Tris pH 7.4, 150 mM NaCl, 5 mM EDTA, protease inhibitor cocktail (Roche# 11836170001), 1 mM Na3VO4, 0.1% SDS). Samples were resolved by 10% SDS–polyacrylamide gel electrophoresis experiments. Blots were incubated overnight at 4 °C with the corresponding primary antibody directed CISH (Cell Signaling Technology #8731) or Akt (Cell Signaling Technology #9272) for Western blotting. Blots were incubated with horseradish peroxidase–conjugated secondary antibodies (Millipore) for 1 hr at room temperature. ECL (enhanced chemiluminescence; SuperSignal West Pico and SuperSignal West Femto, Pierce) was used to visualize protein bands.

### NK cell cytotoxicity assays

Standard 4 hr cytotoxicity assays were completed as described elsewhere^33^. Briefly, splenic NK cells were isolated and suspended in NK cell medium (phenol-red free RPMI 1640 containing 10% FCS, non-essential amino acids, L glutamine and sodium pyruvate, all from Gibco). The indicated target cells were labelled with 15μM Calcein-AM (Life Technologies) for 30 min at 37°C, washed twice and suspended in NK cell medium. Effector and target cells were combined at the indicated ratios in triplicate wells of a round-bottom 96 well plate and incubated at 37°C / 5% CO_2_ for 4 hours. Calcein release was quantified by transferring 100 μL of cell-free supernatant to opaque 96 well plates and measuring fluorescent emission at the appropriate wave-length (excitation filter: 485±9 nm; cutoff: 515 nm; emission: 525±15 nm) using the EnVision Robot Plate Reader.

### Flow cytometry and cell sorting

Single-cell suspensions were stained with the appropriate monoclonal antibody in PBS containing 2% FCS. When necessary, intracellular staining was performed by use of the FoxP3/Transcription Factor Staining Buffer Set (eBioscience) according to the manufacturer’s instructions. LSRII, Fortessa, (BD Biosciences) were used for cell analysis. Antibodies specific for NK1.1 (PK136; 1:100), CD19 (1D3; 1:400), CD3 (17A2; 1:400 or REA641; Miltenyi Biotec; 1:150); CD122 (TM-β1; 1:200), NKp46 (29A1.4; 1:100), KLRG1 (2F1; 1:200), CD27 (SB/199; 1:200), CD11b (M1/70; 1:200), IL-7R (A7R34; eBioscience; 1:200) CD49b (DX5; 1:100), CD49a (Ha31/8; 1:200) Ly49H (3D10; 1:200) Ly49D (4E5; 1:200), NKG2D (C4; 1:200), NKG2A/C/E (20d5; 1:200), Ly49C/I (5e6; 1:100), CD107a (104B; 1:100) and IFN-γ (XMG1.2; 1:100), DNAM-1 (10E5; 1:200); Ki-67 (AF488; 1:50) were from BD Pharmingen unless stated otherwise.

### Cell Counts

123count eBeads (BD Bioscience) beads were added to single cell suspensions prior to flow cytometry. Cell numbers were enumerated according to manufacturers instructions.

### Enumeration of Apoptotic Cells

The enumeration of apoptotic cells was performed using the CellEvent^TM^ Caspase-3/7 Green Flow Cytometry Assay Kit (catalogue #: C10427; Thermo Fisher Scientific) following manufacturer’s instructions.

### Proliferation assay

For assessment of cellular proliferation by CFSE (carboxy fluorescein diacetate succinimidyl diester; Life Technologies #C34554), splenic NK cells were purified from *Cish ^+/+^Ncr1^iCre^* or *Cish^fl/fl^Ncr1^iCre^* mice, then labeled with 0.1 μM CFSE and cultured with various doses of hIL-15 for 5 days, before flow cytometric analysis.

### Cytokine assay

For the measurement of IFN-γ production, splenocytes were activated in 96 well plates with hIL-15 (50 ng/ml), IL-12 (10 ng/ml, Peprotech), IL-18 (100ng/ml), PMA (10ng/ml), Ionomycin (1μg/ml), or coated with anti-NK1.1 (25 μg/ml, BioLegend), anti-NKp46 (10 μg/ml, BioLegend) or NKG2D (10 μg/ml, BioLegend). Cells were incubated with monensin and brefeldin A (BD GolgiPlug and GolgiStop) in complete medium for 4 h at 37°C. The cells were subjected to surface staining and intracellular staining was performed by use of the FoxP3/Transcription Factor Staining Buffer Set (eBioscience).

### Tumor cell lines

B16F10 melanoma cells were obtained from ATCC and were maintained in Dulbecco’s Eagle Modified Medium (DMEM) supplemented with 10% FBS. EO771 cell line was purchased from CH3 BioSystems LLC (Amherst, NY, USA). E0771-GFP^+^-Luciferase were generated as was previously described ^34^ and were maintained in RPMI-1640 media supplemented with 10% FCS. NK-92 were obtained from ATCC were grown in RPMI-1640 (Invitrogen) medium supplemented with 1 mM sodium pyruvate, 100 U/ml penicillin, 100 μg/ml streptomycin and 20% heat-inactivated FCS plus 500 UI of IL-2. The K562 cell line was cultured in RPMI-1640 containing 10% heat-inactivated FCS with 1 mM sodium pyruvate, 100 U/ml penicillin, and 100 μg/ml streptomycin. EBV-LCL cells were a kind gift from R. Childs (NIH) and were cultured in RPMI supplemented with 10% fetal bovine ^35^.

### Experimental tumor metastasis

Single-cell suspensions of 3×10^5^ B16F10 melanoma cells were injected i.v. into the tail vein of the indicated strains of mice. Mice were sacrificed and lungs were harvested on day 14. Lungs from B16F10 injected mice were fixed in PFA 4% overnight to count B16F10 metastases. E0771-GFP^+^-Luciferase breast cancer cells were injected into the tail vein of the indicated strains of mice (5×10^5^ cells/mouse). Luciferase expression was then monitored at day 7 and 14 by bioluminescence using PhotonIMAGER (BiospaceLab), following intraperitoneal injection of luciferin (30 mg/kg). After completion of the analysis organ luminescence was assessed. Orthotopic implantation of breast tumors was performed as previously described ^34^. Briefly, EO771-Luc/GFP cells were suspended in 100 μL of a mixture of PBS/Matrigel (v/v) (Corning). 5 × 10^5^ EO771-Luc/GFP cells were injected into the 4th inguinal mammary fat pads of 6 to 10 week old female C57BL/6 mice. Tumor growth was monitored by caliper measurements and weighted at day 15. Lung and liver were harvested at day 15, Luciferase expression was monitored by bioluminescence using PhotonIMAGER (BiospaceLab), after intraperitoneal injection of luciferin (30 mg/kg). Tumor dissociation was performed as previously described ^34^, the counting of Infiltrated NK cells was performed using countbright absolute counting beads by flow cytometry (ThermoFisher Scientific, #C36950) according to manufacturer’s instructions.

### Sample preparation, RNA sequencing and bioinformatics analysis

RNA isolation from sorted *ex vivo* NK cells was extracted using the RNeasy Plus mini Kit (#74134, QIAGEN, Hilden, Germany), according to the manufacturer’s instructions. Purified RNA was measured using an Agilent 2200 TapeStation System (Agilent) with High Sensitivity (HS) RNA ScreenTapes (#5067-5579, Agilent). For Library construction, Full length cDNA were generated from 4 ng of total RNA using Clontech SMART-Seq v4 Ultra Low Input RNA kit for Sequencing (Takara Bio Europe, Saint Germain en Laye, France) according to manufacturer’s instructions with 9 cycles of PCR for cDNA amplification by Seq-Amp polymerase. Six hundreds pg of pre-amplified cDNA were then used as input for Tn5 transposon tagmentation by the Nextera XT DNA Library Preparation Kit (96 samples) (Illumina, San Diego, CA) followed by 12 cycles of library amplification. Following purification with Agencourt AMPure XP beads (Beckman-Coulter, Villepinte, France), the size and concentration of libraries were assessed by capillary electrophoresis. The library was sequenced on Illumina Hiseq 4000 sequencer as Single-Read 50 base reads following Illuminas’s instruction and base calling were performed using RTA 2.7.7 and bcl2fastq. Approximately 60 million reads per sample were obtained by pooling RNA libraries and performing single-end 50bp sequencing. Sequencing was performed at the GenomEast platform at the IGBMC (Institut de Génétique et de Biologie Moléculaire et Cellulaire).

Single-End reads 35-76bp in length corresponding to *Cish^−/−^* and WT *Cish^+/+^* NK cells (3 biological replicates per group) were quality checked using fastqc ^36^. Low quality bases (Phred quality score less than 30) were filtered out and TrueSeq Adapters were trimmed using trimmomatic ^37^. Reads were mapped to mm10 using subread-align (v1.5.0) ^38^ with default parameters. The aligned reads were summarized at the gene-level using featureCounts^39^, counts were normalized by the size of each library (DESeq2, estimateSizeFactors function) and finally differentially expressed genes (DEG) analysis was performed using DESeq2 package with default parameters ^40^. Genes were considered as DEG if they achieved a false discovery rate of 5% or less. Finally, gene annotation and GO/KEGG pathway enrichment analysis were carried out using Mus musculus (org.Mm.eg.db) AnnotationDbi ^41^ and clusterProfiler (enrichGO and enrichKEGG functions) ^42^ packages from R/Bioconductor.

### Plasmids

pHR-SFFV-KRAB-dCas9-P2A-mCherry ^21^ was purchased from addgene. sgRNAs targeting *Cish* were designed using the broad institute online tool sgRNA design CRISPR(i) (https://portals.broadinstitute.org/gpp/public/analysis-tools/sgrna-design-crisprai). The sgRNA leading to the most important downregulation of CISH protein in western blot was subsequently used in our study (CISH sgRNA sequence: CTGGAGGGAACCAGTGGGCG). This CISH was then inserted into EF1-GFP-U6 vector using EF1-GFP-U6-gRNA linearized SmartNuclease Lentivector Plasmid kit according to manufacturer’s instructions (system biosciences).

### Primary NK cell cultures

Peripheral blood mononuclear cells (PBMCs) were obtained by centrifugation on density gradient of whole blood from healthy volunteers (HV) provided by the local Blood Bank (Etablissement Francais du Sang (EFS). Human NK cells were isolated from peripheral blood mononuclear cells (PBMC) obtained from multiple different healthy volunteers. Negative selection of NK was performed on EasySep™ Human NK Cell Enrichment Kit system (StemCell Technologies). 2.10^6^ sorted NK cells were cultured with 100 Gy-irradiated EBV-LCL cells in a 1:20 ratio, in 20 ml of RPMI supplemented with 10% fetal bovine, 1X GlutaMAX (both from Gibco) & 500 UI/mL IL-2 (Proleukin, Novartis).

### NK-92 & primary NK cells transduction

Transduction was performed as previously described ^22^. Briefly, HEK-293T cells were transfected with plasmids encoding for HIV gagpol (psPAX2), and the viral envelopes, VSV-g or BaEV ^22^ and the plasmid coding for the vector of interest using lipofectamine LTX (Invitrogen). 48hrs later the virus-containing supernatant from HEK-293T cells was concentrated 100-fold using Lenti-X concentrator (Takara). Titration was performed on HEK293T cells (ATCC) using serial vector dilutions. Concentrated vectors were added at the indicated MOI (multiplicity of infection) in presence of retronectin at 10ug/ml (Takara). The plates were then centrifuged at 1,000 g for 1 h and incubated at 37°C during 3hrs. NK-92 or primary NK cells were then added at 1. 10^6^ cells/ml in 500ul of regular medium supplemented with IL-2 500UI in presence of protamine-sulfate at 20ug/ml (Sigma). The plates were then centrifuged at 1,000 g for 1 h and incubated at 37°C overnight. The next day, IL-2-supplemented medium was added to each well. Transduction was assessed by cytometry on day 7 after transduction. NK-92 or primary NK cells were then sorted using the the BD *FACSAria*™ *III* Cell Sorter sorting mCherry and GFP positive cells.

## ACKNOWLEDGEMENTS

This work is supported in France by institutional grants from the Institut National de la Santé et de la Recherche Médicale (Inserm), Centre National de la Recherche Scientifique (CNRS) and Aix-Marseille Université to CRCM; by project grants from the Fondation ARC pour la recherche sur le Cancer (PJA20161204835, PJA20191209406), Groupement des Entreprises Françaises dans la Lutte contre le Cancer (GEFLUC) Marseille-Provence, then the Janssen Horizon Fonds de dotation. P-L.B. is supported by a doctoral fellowship from Ligue Nationale contre le Cancer and V.L. by a doctoral fellowship from the Ministère de l’Education et la Recherche, then from the Fondation ARC pour la recherche sur le Cancer. G.G. was supported by a post-doctoral fellowship from the Fondation ARC pour la recherche sur le Cancer and is currently supported by the Janssen Horizon Fonds de dotation. D.O. is a Senior Scholar of the Institut Universitaire de France. We thank Canceropôle PACA and the Plan Cancer Equipement (#17CQ047-00) for continued support the development of the TrGET preclinical assay platform. The team “Immunity and Cancer” was labeled “Equipe Fondation pour la Recherche Médicale (FRM) DEQ 20180339209” (for D.O.).Centrede recherche en Cancérologie de Marseille (Inserm U1068) and the Paoli Calmettes Institute are members of OPALE Carnot Institute, The Organization for Partnerships in Leukemia, Institut de Recherche Saint-Louis, Hôpital Saint-Louis, 75010 Paris, France. We wish to thank Bernard Malissen and Frédéric Fiore for their expertise to generate Tm1c *Cish^fl/fl^* mice (B6-Cish^tm1cCiphe^) at the Centre d’Immunophénomique (Ciphe) (Marseille, France), Esma Karkeni for participating in the characterization of the *Cish^fl/fl^Ncr1^Ki/+^* mouse strain and Arnaud Capel for excellent animal husbandry. We are grateful to the staff of the staff of the CRCM animal facility for taking care of the mouse strain colonies and the CRCM cytometry platform for FACS analysis.

## AUTHOR CONTRIBUTIONS

P-L.B., S.P., V.L., A.G., E.J., R.C., J.V. and G.G. all performed experiments. G.G., and J.A.N. designed experiments and analysed the data. R.B.D., D.O., E.V., and N.D.H. contributed intellectual input and helped to interpret data. G.G. and J.A.N. led the research program. N.D.H., G.G., and J.A.N. wrote the manuscript.

