## supplemental figures and legends for "Targeting CISH enhances natural cytotoxicity receptor signaling and reduces NK cell exhaustion to improve solid tumor immunity"

Supplementary figures and legend:

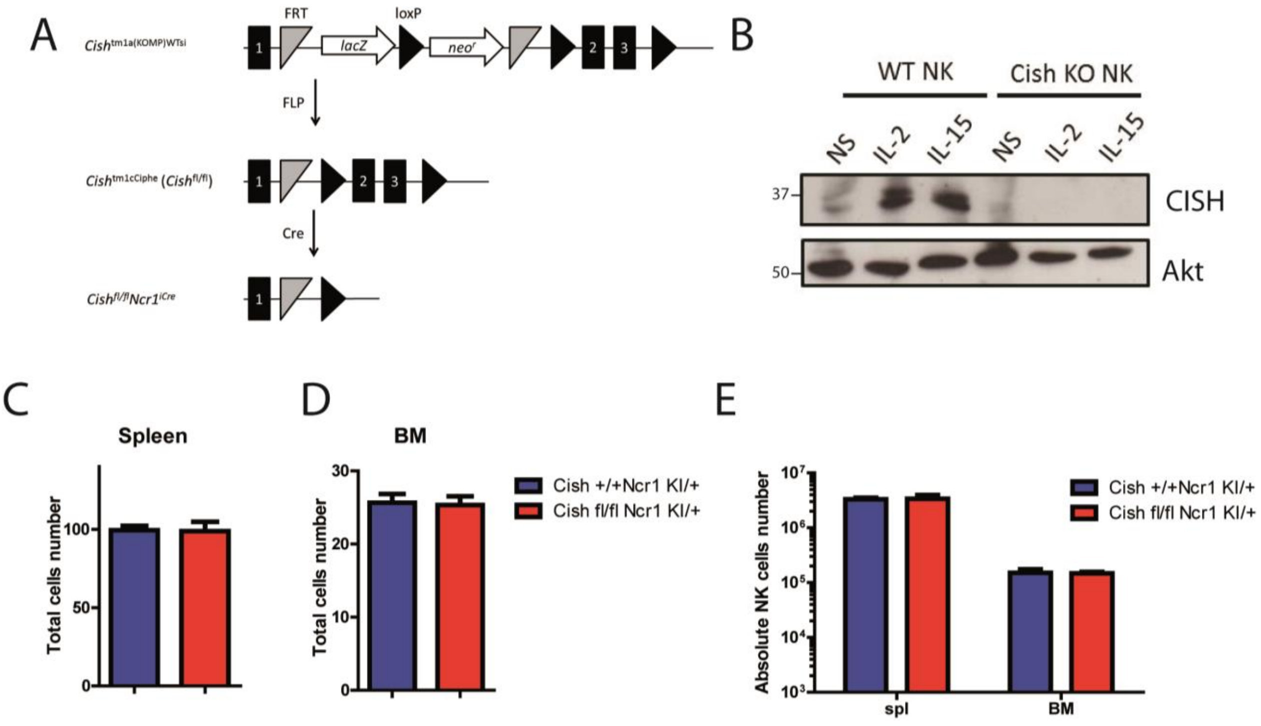


**Supplemental figure 1** (**A**) Conditional *Cish*-deficient mice were generated by breeding mice containing the germline-transmitted Cish$^{tm1a}$ allele to a FLP deleter strain to remove the *lacZ* and *neo$^r$* genes flanked by FRT recognition target sites. LoxP sites

were inserted on either side of exons 2 and 3 to enable Cre-mediated inactivation of the Cish gene in NK cells after breeding to $Ncr1^{iCre}$ mice. (**B**) NK cells were purified from *Cish* $^{+/+}$*Ncr1*$^{Ki/+}$ or *Cish* $^{fl/fl}$*Ncr1*$^{Ki/+}$ spleens, expanded for 6 days with IL-15 and FACS sorted (NK1.1$^{+}$, NKp46$^{+}$, CD3$^{-}$, CD19$^{-}$). After overnight starvation, *Cish* $^{+/+}$*Ncr1*$^{Ki/+}$ and *Cish* $^{fl/fl}$*Ncr1*$^{Ki/+}$ NK cells were either untreated or stimulated for 4 hours with IL-2 or IL-15. Cell lysates were analysed by immunoblotting for Cish or Akt (loading control). (**C**-**D**) Total number of cells purified from *Cish* $^{+/+}$*Ncr1*$^{Ki/+}$ or *Cish* $^{fl/fl}$*Ncr1*$^{Ki/+}$ spleens (**C**) or Bone marrow (**D**). (**E**) Absolute number of NK cells (CD3-, NK1.1+) purified from *Cish* $^{+/+}$*Ncr1*$^{Ki/+}$ or *Cish* $^{fl/fl}$*Ncr1*$^{Ki/+}$ spleens or Bone marrow. (**C**-**E**) n=6 biological replicates mean ± s.e.m.

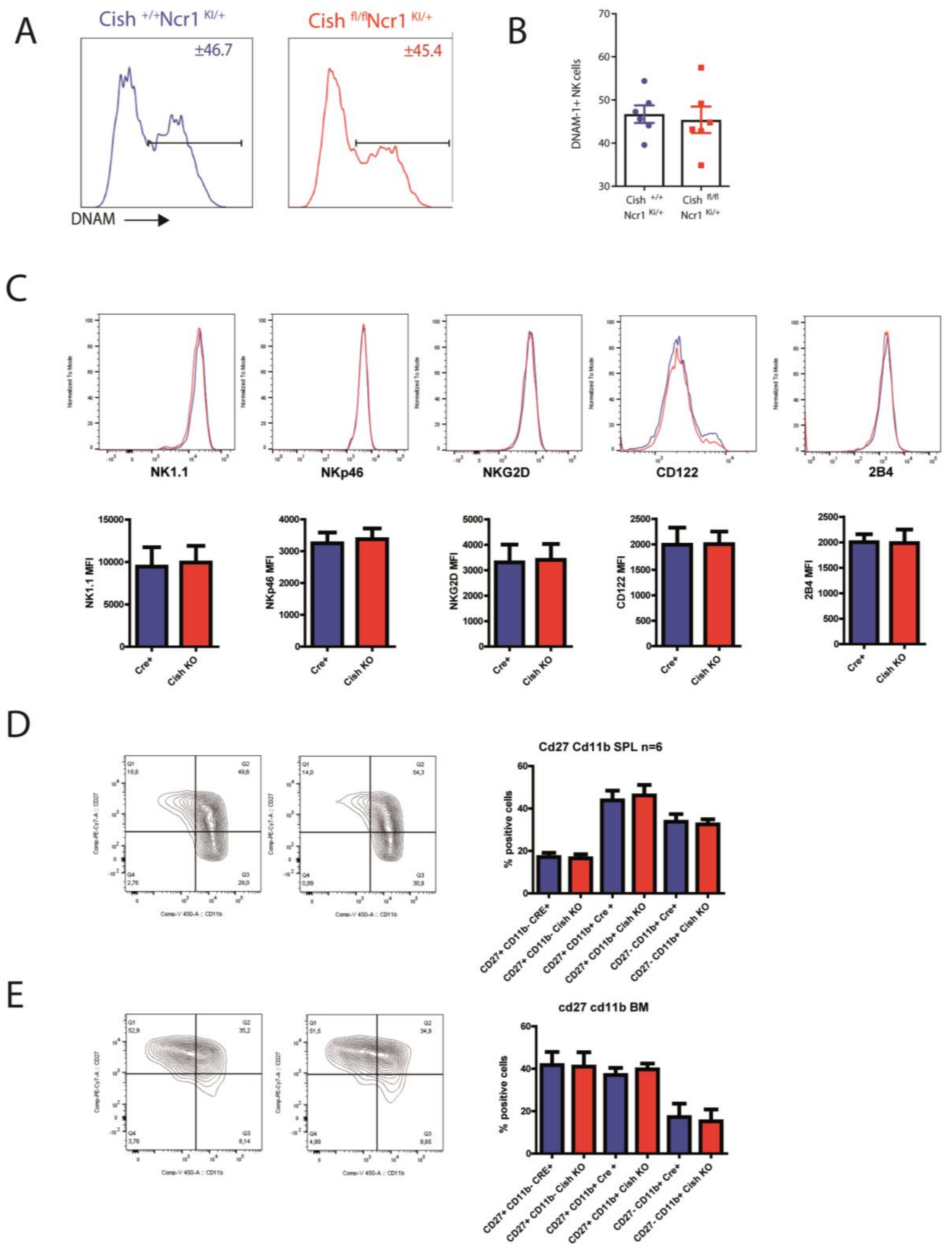
A
Cish +/+Ncr1 KI/+
±46.7
Cish fl/flNcr1 KI/+
±45.4
DNAM
B
DNAM-1+ NK cells
Cish +/+ Ncr1 Ki/+
Cish fl/fl Ncr1 Ki/+
C
NK1.1
NKp46
NKG2D
CD122
2B4
NK1.1 MFI
NKp46 MFI
NKG2D MFI
CD122 MFI
2B4 MFI
Cre+
Cish KO
D
Cd27 Cd11b SPL n=6
% positive cells
CD27+ CD11b- CRE+
CD27+ CD11b- Cish KO
CD27+ CD11b+ Cre +
CD27+ CD11b+ Cish KO
CD27- CD11b+ Cre+
CD27- CD11b+ Cish KO
E
cd27 cd11b BM
% positive cells

**Supplemental figure 2.** Splenic NK cells from *Cish* $^{+/+}$*Ncr1*$^{Ki/+}$ and *Cish* $^{fl/fl}$*Ncr1*$^{Ki/+}$ mice were phenotypically analysed by flow cytometry. (**A-B**) Frequency of DNAM-1$^{+}$ NK cells and representative histogram is showed. (**C**) Representative histogram and Mean Fluorescence intensity (MFI) for NK1.1, NKp46, NKG2D, CD11 and 2B4 receptors are showed. (**D-E**) Frequency of CD27,Cd11b cells were measured within NK cell populations from the spleen, representative FACS plots and histograms are shown. n=6 biological replicates mean ± s.e.m.

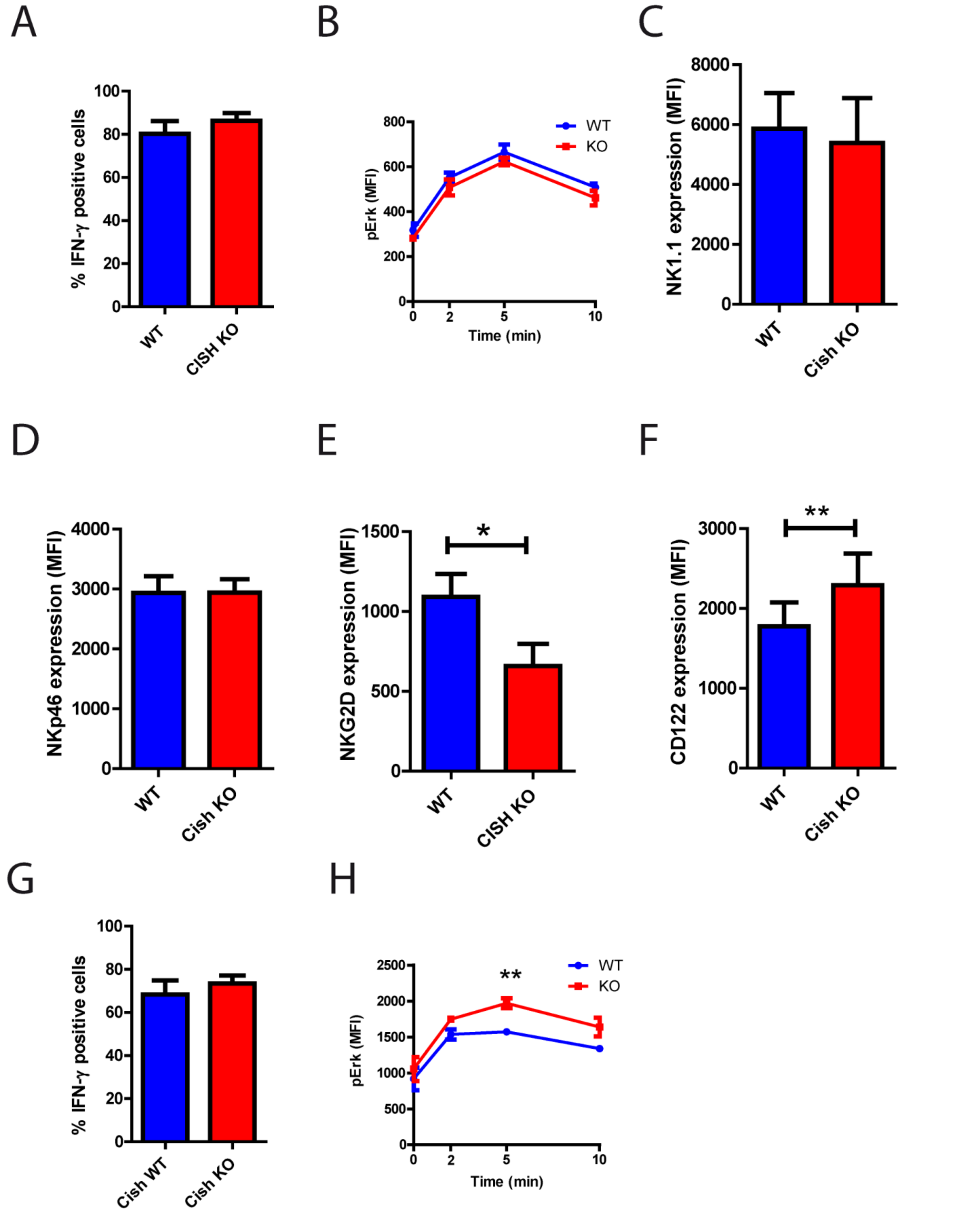


**Supplemental figure 3.** (**A**) splenocytes cells were stimulated with PMA and ionomycin for 4 hrs. Flow cytometric analysis of NK cells and their IFN-γ production was assessed. (**B**) splenocytes cells were stimulated with PMA and ionomycin for 2, 5 and 10 minutes. Flow cytometric analysis of NK cells ERK1/2 phosphorylation was assessed. (**C-H**) Total splenocytes were expanded in IL-2 during 6 days (1000UI/ml), then an overnight IL-2-starvation was performed. (**C-F**) Flow cytometric analysis of NK cells and MFI expression of NK1.1 (**C**), NKp46 (**D**), NKG2D (**E**) and CD122 (**F**) receptors was assessed. (**G**) Expanded splenocytes (LAKs) were stimulated with PMA and ionomycin for 4 hrs. Flow cytometric analysis of NK cells and their IFN-γ production was assessed. (**H**) Expanded splenocytes (LAKs) were stimulated with PMA and ionomycin for 2, 5 and 10 minutes. Flow cytometric analysis of NK cells ERK1/2 phosphorylation was assessed. (**A-F**) n=4 biological replicates mean ± s.e.m. (**G-H**) n=3 biological replicates mean ± s.e.m. *$p<0.05$, **$p<0.01$ (Student's *t*-test).

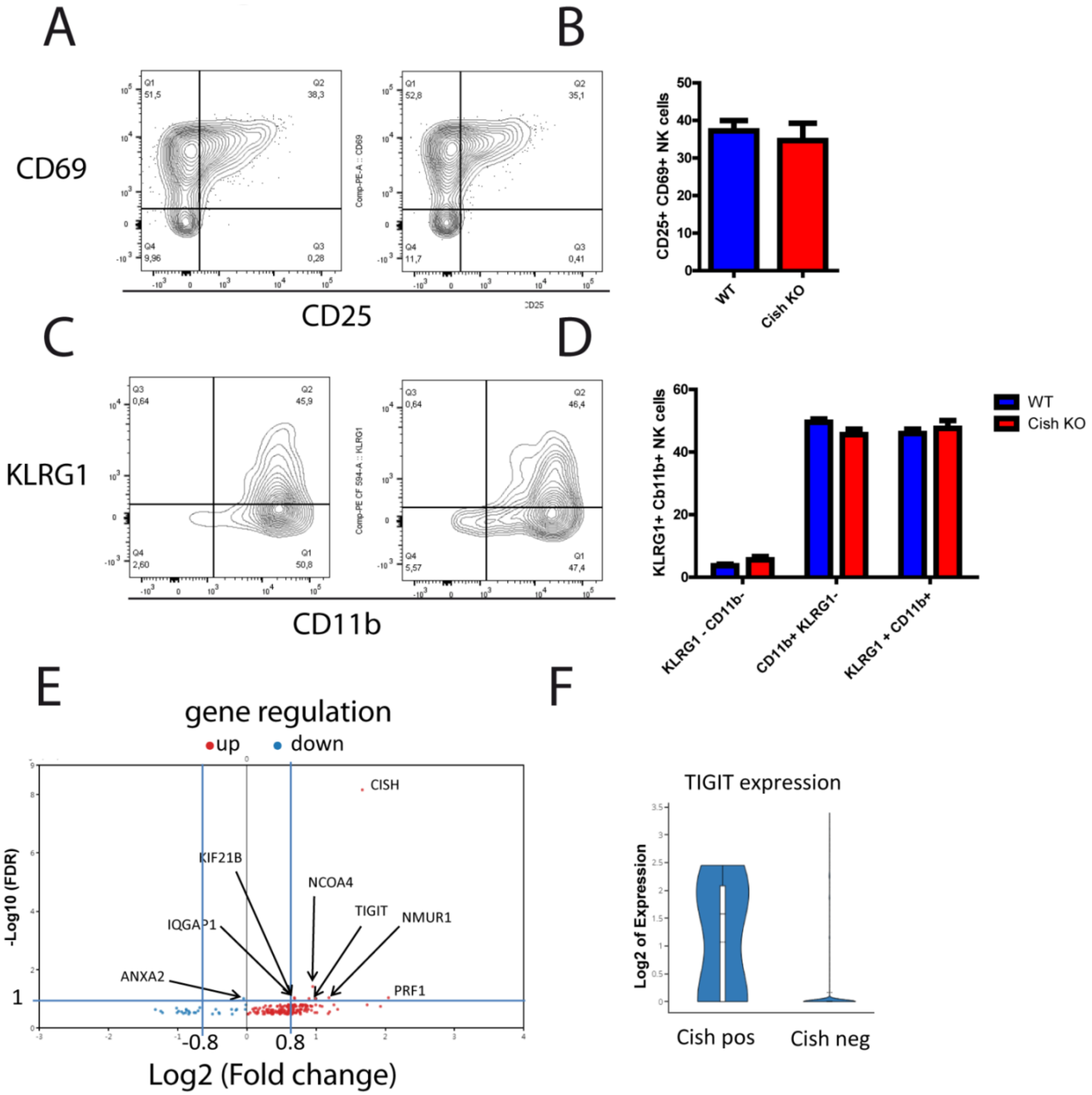


**Supplemental figure 4.** Orthotopic injection of EO771-GFP$^{+}$ Luciferase$^{+}$ breast cancer cells in the mammary fat pad of *Cish* $^{+/+}$*Ncr1*$^{Ki/+}$ and *Cish*$^{fl/fl}$*Ncr1*$^{Ki/+}$ mice. (**A-D**) Flow cytometry was used to quantify frequency of CD25 and CD69 cells (**A-B**) and KLRG1 and CD11b (**C-D**). (E-F). Loupe cell browser software was used to analyse previously published Database of Single cells mapping of immunes cells infiltrated in human breast tumors (Azizi et al., Cell 2018). We isolated infiltrated NK cells (*Cd3-*, *Ncam1+)* then observed the expression of *Tigit* comparing *Cish*+ NK cells with *Cish*- cells. (**E**) A volcano plot of significant (*-log10FDR>1*) differentially expressed genes with a log2fold change above 0.8 or below -0.8 was generated. y-axis is the negative log10 (FDR) value and x-axis is the log2-fold-change of the corresponding gene in *Cish*$^{-/-}$ vs *Cish*$^{+/+}$ comparison. Gene expression: blue = downregulated, red = upregulated. (**F**)Log2 expression of *Tigit* gene *Cish*$^{-/-}$ vs *Cish*$^{+/+}$ infiltrated NK cells.

supll. Figure 5

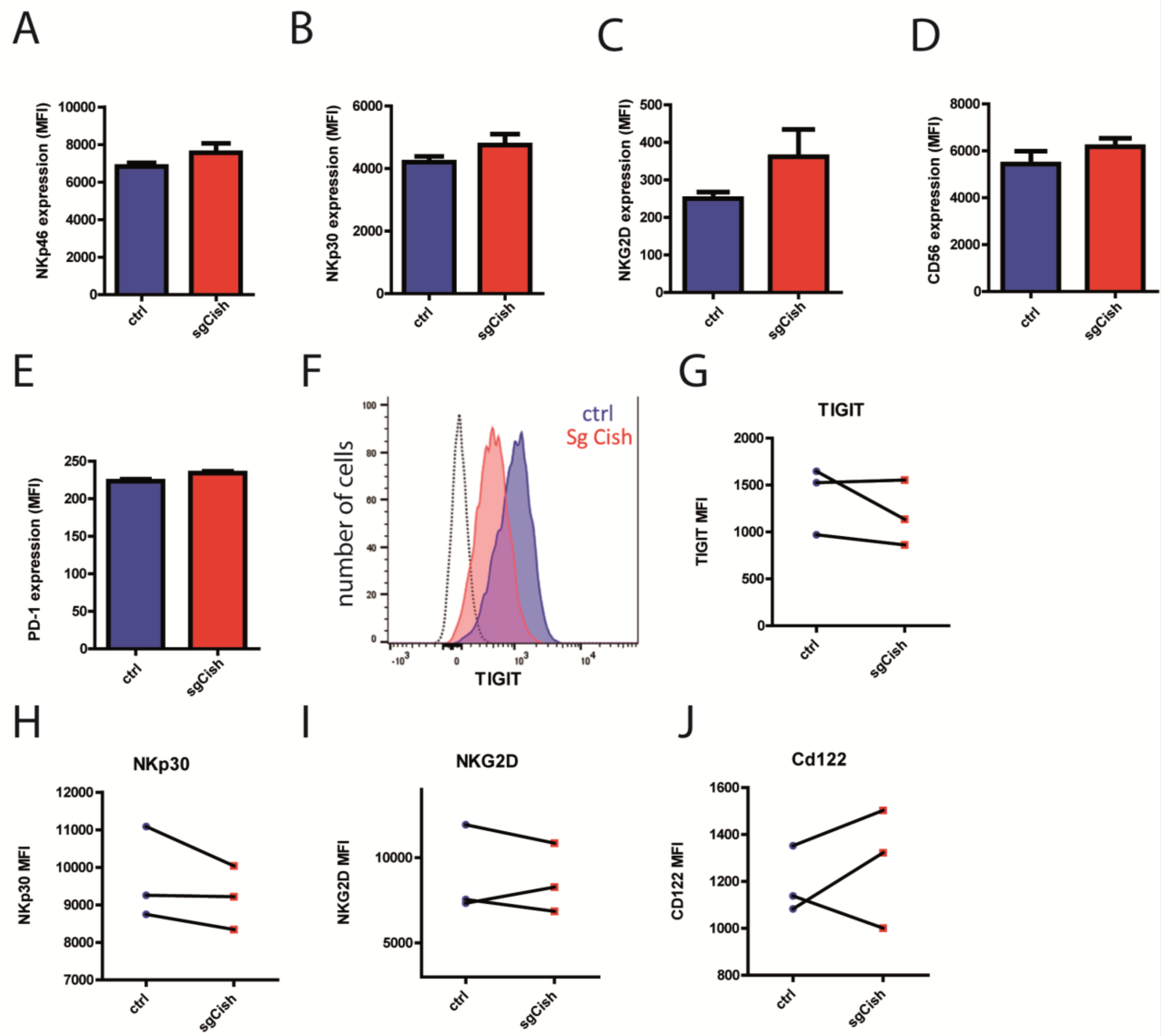


**Supplemental figure 5.** NK-92 / primary NK cells were transduced with a KRAB-dCAS9 construct co-expressing mcherry and a ctrl or Cish SgRNA construct co-expressing GFP. **(A-F)**. Flow cytometric analysis of NK-92 transduced with construct ctrl or sgCISH. MFI expression of NKp46 (**A**), NKp30 (**B**), NKG2D (**C**), CD56 (**D**) and PD-1 (**E**) receptors was assessed. (**F**) representative histogram of TIGIT is showed. (**G-J**) Flow cytometric analysis of primary NK transduced with construct ctrl or sgCISH. MFI expression of TIGIT (**H**), NKp30 (**I**), NKG2D (**J**) and CD122 (**K**) receptors was assessed.
